# Physical and functional interactions of the potassium uptake protein HAK5 and the nitrate transporter NPF6.2 is critical for the mineral nutrition of Arabidopsis

**DOI:** 10.1101/2025.10.03.680249

**Authors:** Laura Morales de los Ríos, Sahar A. Alshareef, Natalia Raddatz, Raúl Carranco, Imelda Mendoza, Thibaut Perez, Claire Corratgé-Faillie, Anna De Luca, Francisco J. Quintero, Magdy M. Mahfouz, Benoît Lacombe, José M. Pardo

## Abstract

Potassium (K^+^) starvation induces the expression of gene *HAK5* encoding a high-affinity K^+^ uptake protein, but how plants perceive the K^+^ status and the signaling intermediaries involved in the response remains largely unknown. To identify key regulators of K^+^ nutrition in Arabidopsis, a genetic screen was performed using an *pHAK5:LUC* reporter line, and a mutant showing stable induction of the reporter under K^+^-sufficient conditions was isolated. Mapping-by-sequencing identified two linked mutations affecting genes involved in K^+^ and nitrate nutrition, namely a loss-of-function in the K^+^ uptake channel AKT1 and a gain-of-function allele of the nitrate transporter NPF6.2/NRT1.4 (NPF6.2^V210M^) that doubled the rate of nitrate transport. We report that the physical interaction of NPF6.2 and HAK5 transport proteins resulted in reciprocal interference. Co-expression in *Xenopus* oocytes of NPF6.2 with the regulatory kinase CIPK23 or the mutant protein NPF6.2^V210M^ alone inhibited HAK5 transport, whereas HAK5 inhibited nitrate transport by NPF6.2 and NPF6.2^V210M^. We conclude that mutation NPF6.2^V210M^ enhanced the nutritional defects associated to the loss of AKT1 function through the inhibition of HAK5. These findings evidence an intimate molecular crosstalk between transporters involved in the mineral nutrition of plants. The mutual interference when both transport systems are operative may represent a novel integrative regulatory mechanism in mineral nutrition.

## INTRODUCTION

Potassium (K^+^) and nitrogen (N) are essential macronutrients for plants, and their uptake and distribution must be coordinated for optimal growth (Raddatz et al., 2020). Potassium is involved in enzyme activation, the control of membrane electrical potential, pH homeostasis, charge balance of inorganic and organic anions, and the regulation of osmotic pressure. Nitrogen, on the other hand, is assimilated into amino acids, proteins, and nucleic acids. Nitrate (NO3^-^) is often the primary nitrogen source in soils and serves as a signaling molecule to the plant (Wang et al., 2018).

The voltage-dependent inward-directed K^+^ channel AKT1 and the high-affinity K^+^-H^+^ symporter HAK5 are the major transport systems for K^+^ nutrition in Arabidopsis (Rubio et al., 2008; Aleman et al., 2011). In K^+^-sufficient conditions (above 200 µM K^+^) AKT1 mediates the energetically downhill entry of K^+^ driven by the electrical membrane potential, whereas under K^+^-limiting conditions *HAK5* is transcriptionally activated. The HAK5 transporter couples K^+^-H^+^ symport and drives K^+^ uptake in the low µM range (Maierhofer et al., 2024). Both systems cooperate at intermediate K^+^ concentrations (Aleman et al., 2011). AKT1 and HAK5 are both activated by the kinase CIPK23 in complex with the ancillary Ca^2+^-binding proteins CBL1/9 (Xu et al., 2006; Ragel et al., 2015; Sanchez-Barrena et al., 2020; Maierhofer et al., 2024). CBL1/9 serve both to activate and to recruit CIPK23 to the plasma membrane. Upon limited K^+^ supply, CIPK23 also enables the release of K^+^ from the vacuolar store through the activation of Two-Pore K^+^ (TPK) channels, in liaison with the tonoplast-bound CBL2 and CBL3 that recruit CIPK23 to the vacuole (Tang et al., 2020).

Nitrate is taken up by an array of dedicated transporters that include Low- and High-Affinity Transport Systems (LATS and HATS). LATS are encoded by the Nitrate Transporter 1 / Peptide transporter gene Family (*NRT1/NPF*) and HATS by the Nitrate Transporter 2 (*NRT2)* gene families. The CIPK23-CBL1/9 module is again a critical regulator of nitrate transporters NPF6.3 and NPF6.2. NPF6.3/NRT1.1 is described as a nitrate ’transceptor’, i.e. a protein having a NO3^-^-sensing function together with the canonical nutrient uptake role (Ho et al., 2009; Bouguyon et al., 2015; Podar and Maathuis, 2022). Signaling of NO3^-^ availability through NPF6.3 elicits the Primary Nitrate Response (PNR), which prepares the plant for NO3^-^ assimilation and alters root architecture and developmental phase transition (Bouguyon et al., 2015; Abualia et al., 2022). NPF6.2/NRT1.4, which is highly similar to NPF6.3, has been linked to NO3^-^ and K^+^ nutrition, and to the gating of NO3^-^ flux through the petiole towards the leaf lamina (Chiu et al., 2004; Morales de los Ríos et al., 2021). The Slow Anion Channels SLAC1 and SLAH3 mediate chloride and nitrate transport in guard cells, while SLAH1, SLAH2 and SLAH3 are engaged in Cl^-^/NO3^-^ acquisition by roots and translocation to the shoot (Cubero-Font et al., 2016; Hedrich and Geiger, 2017). Of these, SLAC1 and SLAH3 are targets of CIPK23-CBL9 (Huang et al., 2023; Maierhofer et al., 2024). Last, the CIPK23-CBL1 module phosphorylates and inactivates the Ammonium Transporters AMT1;1 and AMT1;2 in conditions of low NO3^-^ and high external NH4^+^ that are potentially toxic to the plant (Straub et al., 2017). Together, these data evidence the co-regulation of K^+^ and N-containing nutrients by the master kinase CIPK23, although the fine details of how specific outputs for each nutritional condition are achieved by a single kinase are not fully understood (Ródenas and Vert, 2021).

The acquisition of K^+^ and NO3^-^ is often positively correlated, and this effect has been attributed to improved charge balance during nutrient uptake and long-distance transport (Coskun et al., 2017). Under nutrient-sufficient conditions the root-to-shoot co-transport of K^+^ and NO3^-^ is enhanced, whereas under nutrient-limited conditions the transport of both nutrients is restricted (Lin et al., 2008; Drechsler et al., 2015; Meng et al., 2016). Thus, it appears that plants have developed tools to secure the K/N balance, and not only their absolute contents. However, the precise mechanism(s) underpinning the molecular and biochemical crosstalk between these and other nutrients remain largely unexplored (Raddatz et al., 2020). A known linkage in K/N nutrition is that key transporters for K^+^ and nitrogen-supplying compounds are co-regulated by the same CIPK23-CBL1/9 protein kinase modules (Ródenas and Vert, 2021). Another K^+^/NO3^-^ interaction may occur at the level of NO3^-^ transport proteins. Long-distance transport of K^+^ and NO3^-^ involves xylem loading and unloading. In Arabidopsis, NPF7.3/NRT1.5 in the root pericycle loads NO3^-^ into the xylem vessels, whereas NPF7.2/NRT1.8, which is expressed predominantly in xylem parenchyma cells of roots and aerial tissues, retrieves NO3^-^ from the xylem (Lin et al., 2008; Li et al., 2010). Notably, the antagonistic NO3^-^ fluxes determined by NPF7.3/NRT1.5 and NPF7.2/NRT1.8 are also reflected in their K^+^ profiles (Lin et al., 2008; Drechsler et al., 2015; Meng et al., 2016; Zheng et al., 2016; Li et al., 2017). Moreover, NPF7.3/NRT1.5 has been suggested to operate as a K^+^/H^+^ antiporter that mediates K^+^ release out of root parenchyma cells and facilitate K^+^ loading into the xylem through interaction with transporters AHA2 and/or SKOR (Drechsler et al., 2015; Li et al., 2017; Sena and Kunze, 2023). The NO3^-^ transporter and sensor NPF6.3/NRT1.1 and the closely related protein NPF6.2/NRT1.4 also affect plant growth under K^+^-limiting conditions, and mutants in these proteins showed disturbed K^+^-uptake and root-shoot partition (Fang et al., 2020; Morales de los Ríos et al., 2021). These functional interactions of NPF proteins with K^+^ nutrition have been suggested to be dependent on H^+^-consumption associated with NO3^-^-H^+^ symport, which in turn may affect the operation of pH-sensitive K^+^ channels (Fang et al., 2020; Raddatz et al., 2020).

Here, we expand the molecular links between K^+^ and NO3^-^ nutrition to include the physical and functional interactions between the transport proteins themselves. Through a genetic screen for altered expression of *HAK5*, we found that a gain-of-function mutant of NPF6.2 with greater NO3^-^ transport velocity is a genetic enhancer of the phenotypic maladies associated to a dysfunctional *akt1* allele, a background in which HAK5 becomes the critical K^+^-uptake system. We show that protein HAK5 interacted physically with NPF6.2 and the nitrate transceptor NPF6.3, and the interaction was responsive to K^+^ supplementation. Surprisingly, these physical interactions resulted in mutual functional interference between K^+^ and NO3^-^ transporters in *Xenopus* oocytes, especially when co-expressed with the common regulators CIPK23-CBL1. These results demonstrate a novel and intimate molecular crosstalk between transporters involved in K^+^ and NO3^-^ nutrition. We present a model in which mutual interference when both transport systems are operative owing to a nutritional imbalance of low K^+^/NO3^-^ ratio may represent a novel integrative regulatory mechanism in mineral nutrition.

## RESULTS

### Mutagenesis of reporter line HAK5:LUC

An EMS-mutagenized population of the Arabidopsis line expressing the chimeric reporter gene *pHAK5:LUC* (Jung et al., 2009) was screened for altered expression of luciferase under low (0.1 mM) and high (5 mM) K^+^ availability conditions (Supplemental Figure S1). One mutant line (L14) that consistently showed enhanced luminescence in the range of 1-5 mM K^+^ was selected for further analyses (Figure 1). Enhanced luminescence was stable for at least 7 generations, and manifested also in regular ½MS medium containing 10 mM K^+^, 19.7 mM NO3^-^ and 10.3 mM NH4^+^ (Supplemental Figure S2A and B). A diagnostic PCR and sequencing of the *pHAK5:LUC* amplicon confirmed that the reporter gene remained unmodified by the mutagenesis. Two congenic sublines showing a strong mutant phenotype, L14-1 and L14-3, were selected to analyze their response to K^+^ supplementation in LAK medium, which was designed to be nominally free of K^+^ and Na^+^ (Barragan et al., 2012). Compared to the parental line, L14 plants showed stunted vegetative growth and developmental defects, including a more branched floral stalk and early flowering time (Figure 2 and Supplemental Figure S3). Importantly, L14 plants had reduced K^+^ content in LAK medium with 1 mM K^+^ (Figure 2), which explained the transcriptional induction of the reporter gene.

**Figure 1.**
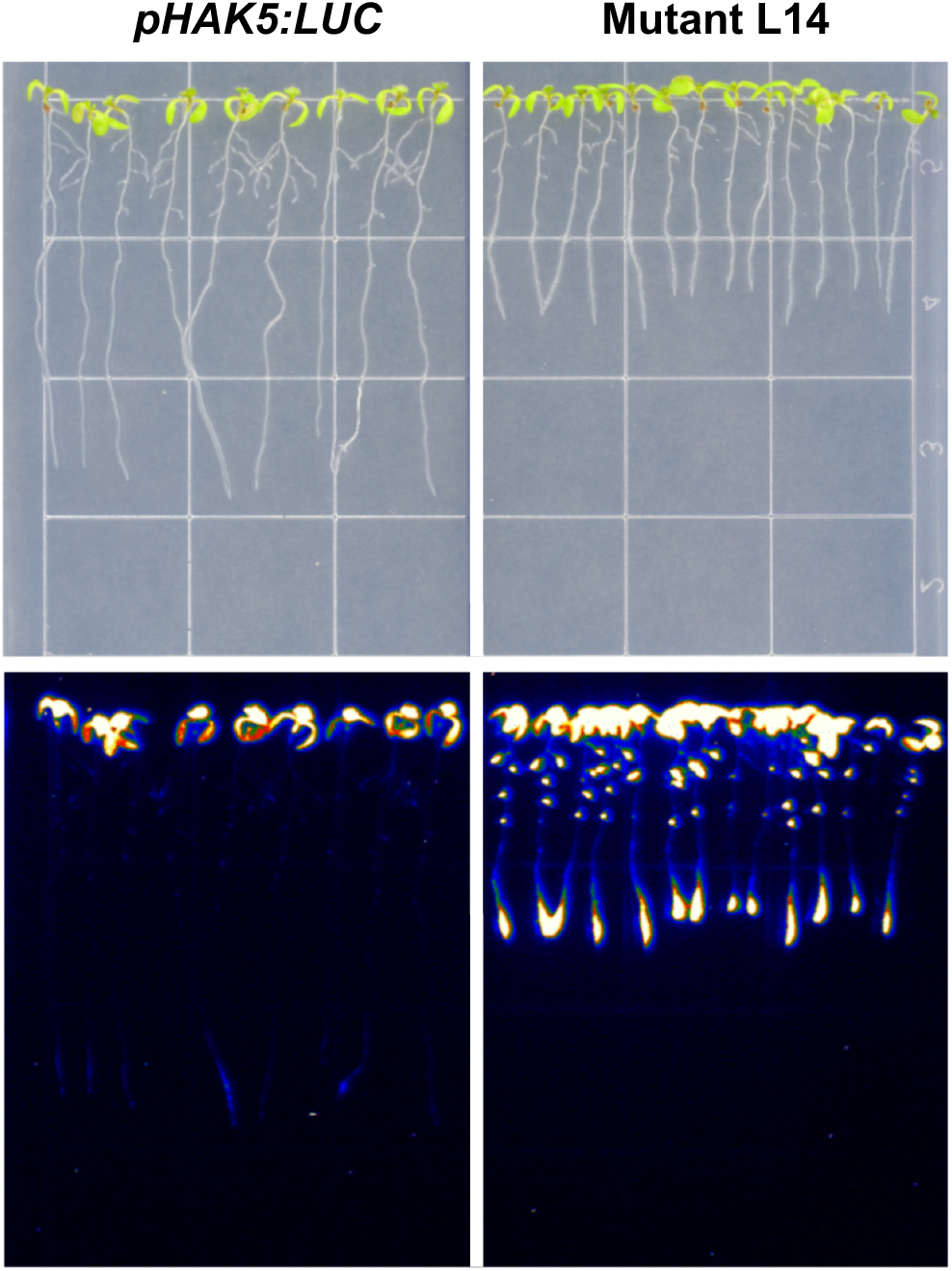
Screening of EMS-mutagenized *pHAK5:LUC* seedlings by luminescence. Four-day old L14 seedlings germinated on basal medium were transfered to 5 mM KCl medium, and grew for additional 3 days. Stronger luminescence signals were detected in L14 mutant seedlings compared to the parental line *pHAK5:LUC*.

**Figure 2.**
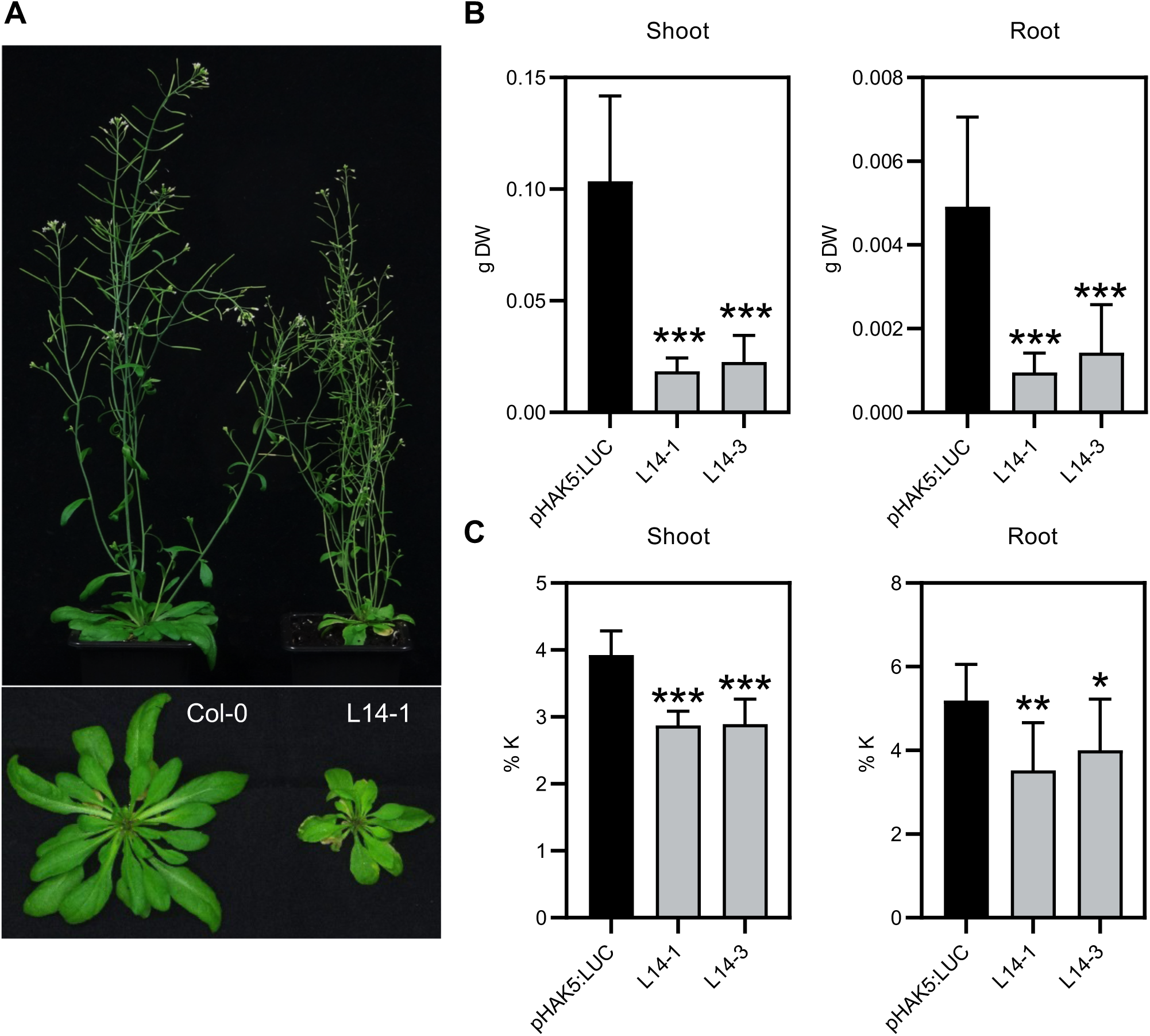
Impaired development and K^+^ content of mutant L14. **(A)** WT and L14 plants grown for 30 days on soil in a controlled growth chamber with the day/night regime of 16/8 hours, 23/20 °C, 70% relative humidity, and 120–140 μmol m^-2^ s^-1^ PAR. **(B,C)** Dry weight (B) and K^+^ content (C) of L14-1 and L14-3 plants grown in hydroponic culture in LAK 1 mM K medium for 30 days. Statistically significant differences between mutant sublines and the parental *pHAK5:LUC* are indicated (Student’s t-test. \**p* <0.1 \*\**p* <0.05. \*\*\**p* <0.01).

Next, plants of the M5 generation were backcrossed to the parental line *pHAK5:LUC* to obtain BC1F1 plants homozygous for the reporter gene and heterozygous for the underlying mutations. These seedlings showed a semidominant luminescence phenotype compared to the parental and L14 lines (Supplemental Figure S2C). Quantification of radiance at the root tips were 0.32 ± 0.07 P·cm^-2^·SR^-1^ (n=7) for L14, 0.22 ± 0.06 (n=7) for BC1F1 seedlings, and background levels for the *pHAK5:LUC* control. Selfing of BC1F1 plants produced a BC1F2 generation segregating 3:1 for enhanced luminescence, indicative of a monogenic mutation or closely linked mutations. To identify the underlying mutation(s) in L14 plants, segregating BC1F2 seedlings having the mutant phenotype or not were bulked (n=20) and submitted to genome re-sequencing. Results showed high-frequency single nucleotide polymorphisms (SNPs) affecting 24 genes in mutant plants, all closely linked in chromosome 2 (Supplemental Table S1). Eight loci showed a bulk 100% frequency of mutant SNPs, whereas the bulk frequency of WT alleles ranged from none to 56.5%. These results are coherent with a mixed population of homozygous and heterozygous mutants within the bulked mutant population caused by the semidominant nature of the underlying mutation(s).

Among the genes having 100% frequency of mutant SNPs was At2g26650, encoding the voltage-dependent K^+^-selective channel AKT1 that has a prevalent function in K^+^ capture by roots (Hirsch et al., 1998). The mutation in L14 plants introduced a premature stop codon at residue 135 of AKT1 (AKT1^W135Z^) predicted to yield a truncated protein containing only the first and second transmembrane segments. To determine whether this truncation produced a loss-of-function (LoF) allele of *AKT1*, the mutant and wild-type channel proteins were expressed in *Xenopus* oocytes to have their transport activity measured by the Two-Electrode Voltage-Clamp technique (TEVC). Samples injected with cRNA encoding AKT1 and the activator kinase complex CIPK23/CBL1 showed robust inward-directed K^+^ currents (6 μA at -180 mV), whereas oocytes expressing AKT1^W135Z^ and CIPK23/CBL1 had background currents (1 µA at -180 mV) similar to water-injected oocytes (Figure 3A,B). Because a functional AKT1 channel is a tetrameric protein whose activity can be modified by the interaction with the KC1 channel subunit (Duby et al., 2008), we also tested whether AKT1^W135Z^ could have a dominant negative effect on the wild-type AKT1 channel, thereby explaining the semidominant nature of mutation(s) in L14 plants. However, co-expression of AKT1^W135Z^ with AKT1 and CIPK23/CBL1 produced currents of similar magnitude to these of AKT1 and CIPK23/CBL1 (Figure 3C). Similar results were obtained upon co-expression of AKT1^W135Z^ and AKT1 in yeast cells impaired in the endogenous system for K^+^ uptake. AKT1 restored growth in low-K^+^ media equally well with and without the co-expression of AKT1^W135Z^ (Figure 3D). Together, these results demonstrate that the truncated protein AKT1^W135Z^ is inactive for K^+^ transport and does not inhibit wild-type AKT1, implying that AKT1^W135Z^ is not likely to interfere with another voltage-gated K^+^ channel of the *Shaker-*like family (Jegla et al., 2018).

**Figure 3.**
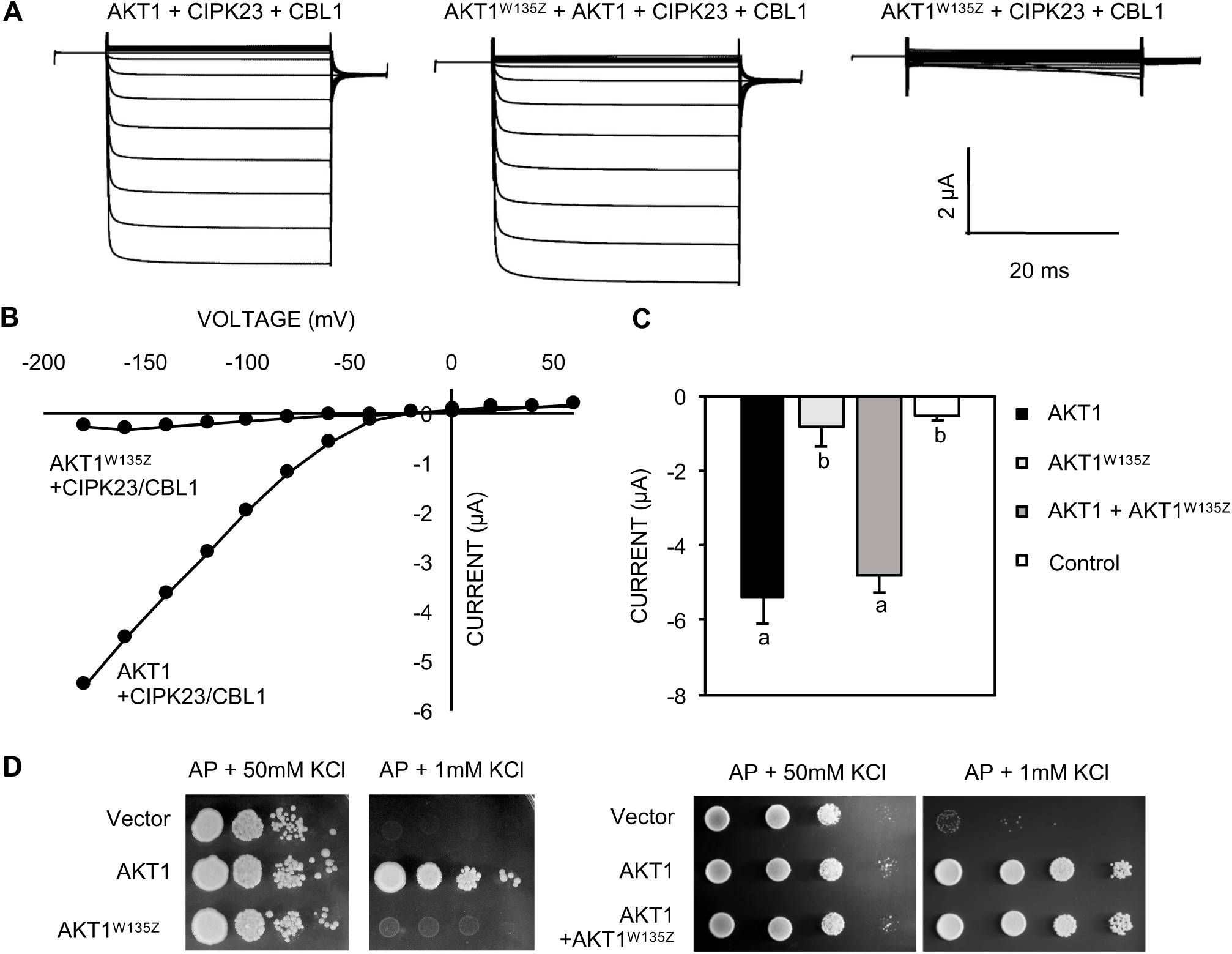
Mutant AKT1^W135Z^ is an inactive channel. **(A)** Currents mesured by TEVC in *Xenopus* oocytes expressing CIPK23, CBL1 and the canonical AKT1 channel and/or AKT1^W135Z^ in the presence of 100 mM KCl in response to a ramp of membrane potentials from -180 to +60 mV. **(B)** Steady-state currents-voltage curve corresponding to currents shown in (A). **(C)** Instantaneous currents collected at -180 mV in oocytes expressing CIPK23/CBL1 together with AKT1 (n=5), AKT1^W135Z^ (n=4), AKT1 + AKT1^W135Z^ (n=4) or water-injected control (n=2). Letters show statistical differences between groups at a significance level of p<0.05 by ANOVA (Tukey’s test). **(D)** Potassium uptake in *S. cerevisiae*. Cells of strain TTE9.3 (*trk1 trk2 ena1-4*) were transformed with the empty vector or the cDNAs encoding AKT1^W135Z^ and/or AKT1. Yeast transformants were grown overnight in AP medium with 50 mM KCl. Cells were washed and 5 µL of serial decimal dilutions spotted onto plates of AP medium supplemented with the indicated KCl concentration. Plates were incubated at 28°C for 3–4 days.

HAK5 and AKT1 act coordinately to enable plant growth in a wide range of external K^+^ concentrations (Gierth et al., 2005; Rubio et al., 2008; Aleman et al., 2011). The above results suggested that the loss of AKT1 was causing the ectopic activation of HAK5 to restore K^+^ uptake. Therefore, we next compared the behavior of L14 plants to that of the knock-out mutant *akt1-2* (Rubio et al., 2008). Wild-type, *akt1-2*, and L14 plants were grown in hydroponic culture with LAK medium with 0.1 and 1 mM KCl in long-day conditions for 30 days. The L14 plants were much smaller than *akt1-2*, and their root K^+^ content was significantly reduced, beyond the decline induced by the *akt1-2* mutation compared to the control. By contrast, the shoot K^+^ content of *akt1-2* and L14 were similarly reduced compared to the wild-type. The water content of L14 plants was also lower than that of wild-type and *akt1-2* controls (Figure 4A-C).

**Figure 4.**
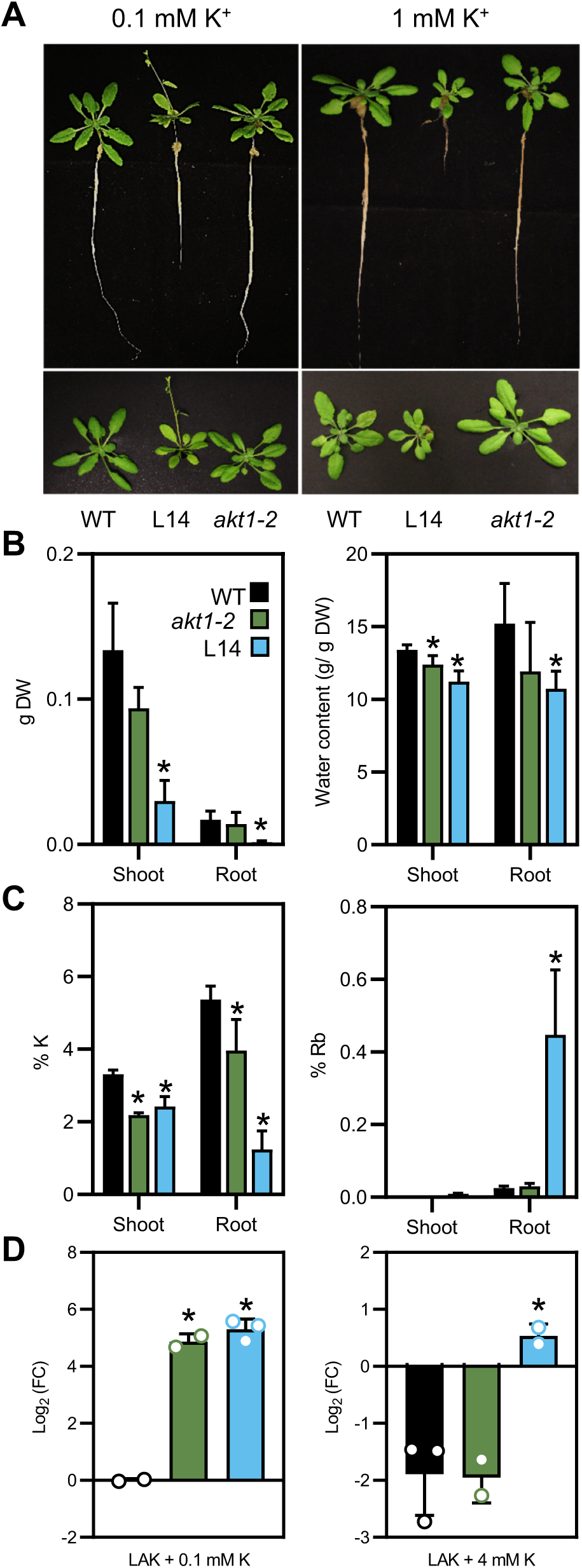
(new). Comparison of *akt1-2* and L14 lines. **(A)** Plants grown for 30 days in hydroponic LAK medium with 0.1 and 1 mM K^+^. **(B)** Dry weight and water contents of 30-day old plants grown in LAK with 1 mM K^+^. Statistical differences of mutant lines relative to the WT were determined by ANOVA (Tukey’s test), * indicates *p*<0.05. **(C)** Rb^+^ uptake. Plants grown for 30 days in LAK with 1 mM K^+^ were deprived of K^+^ for 3 days and transferred to medium containing 50 µM RbCl for 1 hour. Bars show the mean ± SD (n=4 plants) of K^+^ and Rb^+^ contents. Statistical differences were determined by ANOVA (Tukey’s test), * indicates *p*<0.05. **(D)** Expression of the endogenous *HAK5* gene in lines *pHAK5:LUC, akt1-2* and L14. Transcript abundance was quantified in roots of plants grown for 30 days in LAK medium supplemented with 0.1 or 4 mM KCl. *UBQ10* was used as reference gene and *HAK5* expression in line *HAK5:LUC* at 0.1 mM KCl was used for data normalization. * means *p*<0.05 by Tukey’s test, n=3. The experiment was repeated twice with similar results.

Rubidium (Rb^+^) uptake is often used as a proxy to short-term K^+^ uptake by plants and it allows monitoring the high-affinity K^+^ uptake elicited by K^+^ starvation, a process that reflects the activation of HAK5-dependent transport. Hence, we tested Rb^+^ uptake in plants before and after K^+^ starvation (3 days in LAK without supplemental K^+^). In starved plants, Rb^+^ uptake in the high-affinity range (50 µM RbCl) was largely enhanced in L14 compared to *akt1-2* and wild-type plants (Figure 4C). Notably, non-starved L14 plants grown in LAK with 1 mM K^+^ already showed enhanced Rb^+^ uptake at 50 µM despite having K^+^ contents comparable to that of *akt1-2* plants (Supplemental Figure S4). Accordingly, the mRNA levels of the endogenous *HAK5* gene was greater in L14 plants compared to those of parental and *akt1-2* lines in plants grown in 4 mM KCl (Figure 4D). Taken together these results demonstrate that the L14 phenotype is more extreme than that of plants lacking only the AKT1 channel, which together with the semidominant nature of the genetic lesion, strongly implied than L14 contained at least another mutation besides AKT1^W135Z^ giving rise to the strong phenotype observed.

### Mutation in gene NPF6.2/NRT1.4 contributes to the L14 phenotype

Contiguous to the *AKT1* locus is the *NPF6.2/NRT1.4* gene (At2g26690). The frequency of mutant and wild-type SNPs matched those of *AKT1* (Supplemental Table S1) indicating a strong gene linkage. *NPF6.2* encodes a nitrate transporter involved in NO3^-^ transport that also impinges on K^+^ nutrition (Chiu et al., 2004; Morales de los Ríos et al., 2021). The SNP in this locus predicted the change of residue Val210 to methionine in the NPF6.2 protein (NPF6.2^V210M^). The presence of this SNP in line L14 was confirmed by genomic PCR and sequencing of the amplicon.

NPF6.2/NRT1.4 is a plasma membrane protein that mediates NO3^-^ transport in *Xenopus* oocytes and yeast (Chiu et al., 2004; Morales de los Ríos et al., 2021). To analyze the consequences of mutation V210M on the activity of NPF6.2, wild-type and mutant proteins were expressed in *Xenopus* oocytes and incubated for two hours with ^15^N-labeled NO3^-^, after which the ^14^N/^15^N ratio was determined in dried oocytes. Measurement of ^15^NO3^-^ uptake from external concentrations ranging from 0.5 to 30 mM KNO3 showed a low-affinity uptake kinetics in which NPF6.2^V210M^ doubled the accumulation rates of the wild-type protein (Figure 5A). ^15^NO3^-^ uptake by NPF6.2 protein variants was not stimulated by either acidic pH or increased concentrations of K^+^ (Figure 5B,C). NPF6.2 transport was also analyzed by mutant complementation in the nitrate-utilizing yeast *Hansenula polymorpha* (Morales de los Ríos et al., 2021). When expressed in the *ynt1* mutant unable to take NO3^-^ from the media, the NPF6.2^V210M^ mutant protein afforded more robust growth than the wild-type protein in selective medium with 0.5 mM NO3^-^. The difference was cancelled in control medium with 5 mM NH4^+^ as the nitrogen source (Figure 5D). Together, these data indicate that V210M is a gain-of-function (GoF) mutation producing enhanced transport rates in NPF6.2, which is coherent with the semidominant behavior of the genetic lesion in L14 plants.

**Figure 5.**
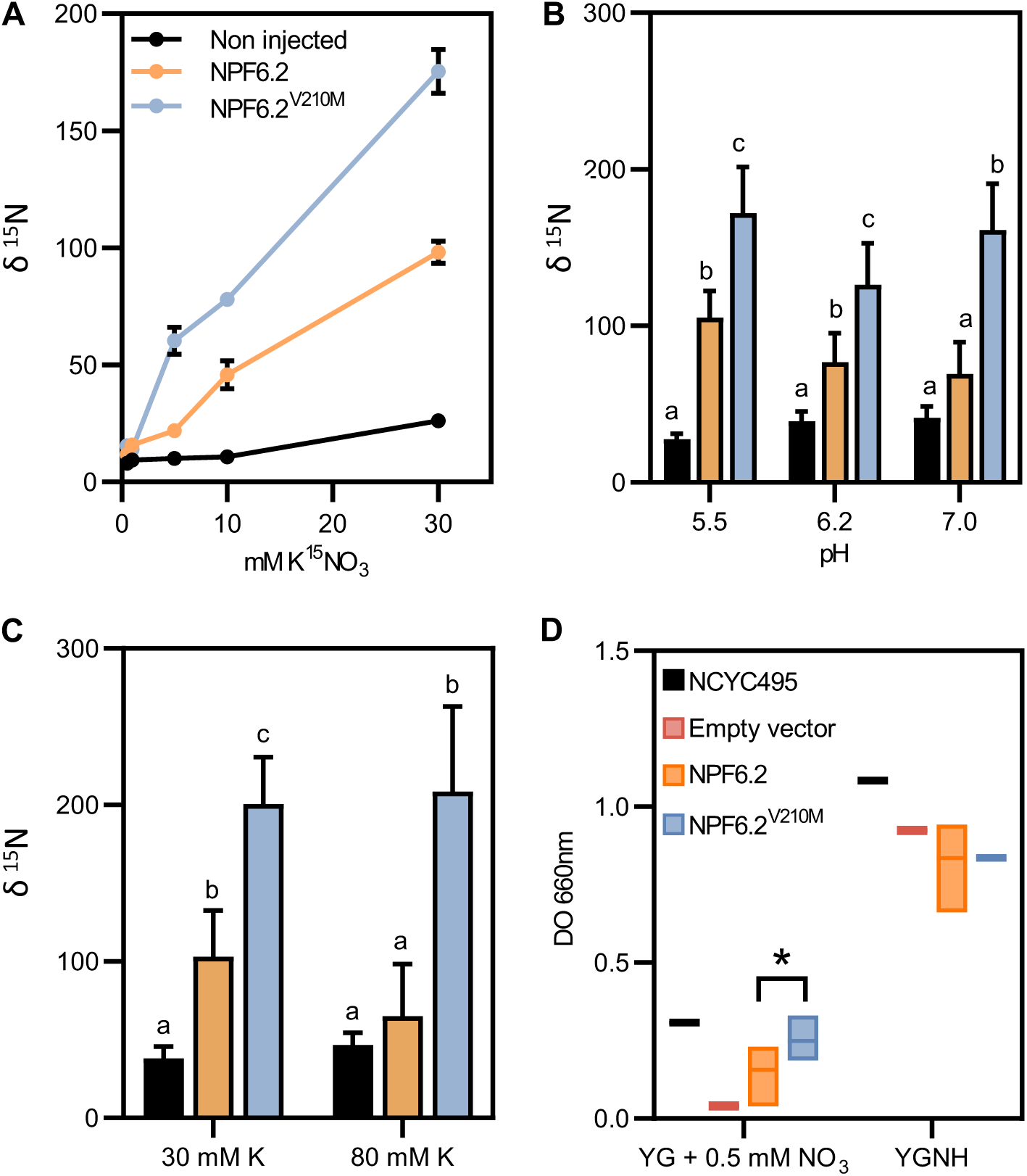
Nitrate transport by NPF6.2 and NPF6.2^V210M^ in heterologous systems. **(A)** ^15^N-nitrate accumulation in oocytes incubated in ND96 medium with varying concentrations of ^15^N-labelled KNO_3_ at pH 5.5. The ^14^N/^15^N ratio was determined in dried oocytes. Data are mean and SD (n=5-10 oocytes). **(B)** ^15^N-nitrate accumulation with 30 mM of KNO_3_ and different pH. Data are mean and SD (n=5-10 oocytes). Letters indicate significant differences (*p*<0.05) by ANOVA with Tukey’s post-hoc test. **(C)** ^15^N-nitrate accumulation in the presence of 30 mM KNO_3_ and 30 or 80 mM KCl, pH 5.5. Data are mean and SD (n=5–10 oocytes). Letters indicate significant differences (*p*< 0.05) by ANOVA with Tukey’s post-hoc test. **(D)** Expression in the yeast *Hansenula polymorpha*. Single colonies of the parental strain NCYC495 (*YNT1*), and of the *Δynt1* mutant transformed with the empty vector, NPF6.2 or NPF6.2^V210M^ were grown at 37°C in YGNH medium. Cultures were washed when reached OD660nm ∼ 1, diluted 1:1000 and incubated for 48 hours at 37°C in YG medium with 0.5 mM KNO_3_ or YGNH control medium. Box histograms frame the maximum and minimum values of OD660 ±SD (n=5) and the central horizontal line represents the mean of the values. Student’s *t*-test for statistical differences between NPF6.2 and NPF6.2^V210M^ (* means *p*<0.1).

We next compared L14 and *akt1-2* plants with the loss-of-function mutant *npf6.2* (a.k.a. *nrt1.4-2*) (Chiu et al., 2004). The reduced petiole NO3^-^ content that had been described for *npf6.2* plants was confirmed, whereas both L14 and *akt1-2* plants had wild-type NO3^-^ levels in petioles (Supplemental Figure S5). L14 plants manifested a greatly compromised growth compared to *akt1-2* and *npf6.2* single mutants (Supplemental Figure 6A). However, L14 and *npf6.2* plants had a similar rate of ^15^NO3^-^ uptake at 0.5 and 5 mM K^15^NO3 external concentration (Supplemental Figure 6B). This implies a minor role of NPF6.2 in net NO3^-^ acquisition since both GoF and LoF mutations had little effect in total ^15^N labeling. To analyze the interplay between nutrient status and the genetic makeup of L14 plants, the NO3^-^ and K^+^ contents in rosette and roots of *akt1-2*, *npf6.2* and L14 plants grown hydroponically in LAK medium modified to give 2 mM K^+^ and 1 mM NO3^-^ final concentrations (see Methods). After 3 weeks, plants were transferred to the fresh medium with 0.5 or 5 mM KNO3 for treatments with low- and high-K^+^/NO3^-^, respectively. In 5 mM KNO3, both *akt1-2* and L14 plants had reduced root K^+^ contents but wild-type levels in the shoot. L14 had a somehow greater K^+^ shoot/(shoot+root) ratio compared to the other lines (WT *vs* L14, *p*=0.142) (Supplemental Figure 6E). When the KNO3 concentration was reduced to 0.5 mM, the *akt1* and L14 shoots showed reduced K^+^ content, and the high K^+^ shoot/(shoot+root) ratio trend in L14 plants was exacerbated owing to the collapse of the root K^+^ content (Supplemental Figure 6C). For nitrate, the content appeared to be mainly dependent on the KNO3 dose with relatively little contribution of the genetic background. At 5 mM KNO3 there were no significant differences between lines (Supplemental Figure 6F). At 0.5 mM KNO3 both *akt1-2* and L14 plants, having in common a defective *akt1* gene, displayed a moderately altered NO3^-^ shoot/(shoot+root) ratio (Supplemental Figure 6D). Taken together, these data supported that the anomalous K^+^ distribution in L14 plants was a consequence of the GoF mutation NPF6.2^V210M^ combined with a LoF mutation in *AKT1*, whereas mutations in L14 had little effect on NO3^-^ uptake and distribution.

### Additional mutations in line L14

The presence of SNPs with 100% frequency in genes *VPS2.1 (AT2G06530), PMT1 (AT2G16120), GGPPS4 (AT2G23800), FUC1 (AT2G28100), GLR2.7 (AT2G9120), PARP1 (AT2G31320), CDF4 (AT2G34140), ERH1 (AT2G37940)* in the L14 line was confirmed by genomic PCR and sequencing of the amplicons (Supplemental Table S1). To purge the additional mutant alleles detected in the mapping population we conducted a series of experiments to evaluate their possible contribution to the phenotype of the L14 line. Mutation in gene At2g24755 was not pursued further because it encoded a protein of unknown function in which no amino acid change was predicted resulting from the SNP.

The SNP detected in gene *VPS2.1* causes the amino acid replacement S4L. VPS2 participates in the ESCRT-III complex *(Endosomal Sorting Complex Required for Transport)* and loss-of-function of VPS2.1 is embryo lethal (Katsiarimpa et al., 2011). Hence, it is unlikely that the S4L mutation suppressed VPS2.1 activity.

Mutation in gene *PMT1* (*Polyol/Monosaccharide Transporter 1*) causes the amino acid change G299E. *PMT1* is separated by an intergenic region of 6602 nucleotides from its homologue *PMT2*. Both genes share 93.6% amino acid sequence identity, and are possibly the result of gene duplication. In addition, PMT1 and PMT2 have similar functional properties and cellular localization (Klepek et al., 2010). The phenotype of knock-out mutant for PMT1 has not been described, but given the degree of homology with PMT2, their similar function and expression pattern, it would be expected that a deficiency of one of the two transporters to be compensated by its homologue.

The mutation in gene *GGPPS4* results in the G250E change. Protein GGPPS4 is located in the endoplasmic reticulum of floral and root procambium cells (Okada et al., 2000), where it catalyzes the formation of geranyl-geranyl diphosphate in the mevalonic acid pathway. Loss-of-function mutants for GGPPS4 and other proteins in this pathway have low embryo viability. In addition, GGPPS4^G250E^ does not belong to critical domains for substrate-specific or catalytics (Beck et al., 2013; Kopcsayová and Vranová, 2019).

FUC1 (α1,3-fucosidase) is implicated in primary cell wall stability. Mutants deficient for this protein do not manifest any developmental defects, despite accumulating the enzyme substrate (Kato et al., 2018). Thus, its involvement in the L14 phenotype could be discarded.

Gene *GLR2.7* encodes a transporter of the glutamate receptor-like channel family (Lacombe et al., 2001). Although the role of glutamate in plants is poorly understood, it has been shown to depolarize the plasma membrane and increase the concentration of calcium in the cytosol (Dennison and Spalding, 2000; Meyerhoff et al., 2005). The SNP detected in gene GLR2.7 causes an exchange C/T in the first intron that did not affect *GLR2.7* gene expression (Supplementary Figure S7A). This made unlikely that the SNP contributed to the L14 phenotype.

Mutation in gene *PARP1* gene causes the change R391H. PARP1 protein, like its homologue PARP2, is an ADP-ribose polymerase involved in DNA repair and implicated in abiotic stress response (Vanderauwera et al., 2007). However, single and double mutants in PARPs do not show phenotypes under stress conditions (Rissel et al., 2017). This may be due to a functional redundancy of the two PARP proteins, as well as the action of other proteins with similar function. Therefore, we could expect that even if the PARP1^R391H^ mutation caused an alteration in protein function, this would be compensated by functional redundancy.

The transcription factor CDF4 promotes differentiation of root columella cells (Pi et al., 2015). To confirm whether the mutation in *CDF4* affects root differentiation in the L14 mutant, its root structure was analyzed by propidium iodide staining. The apical root of WT and L14 lines showed a properly organized and differentiated quiescent center (QC) and columella (C) without significant differences between lines (Supplementary Figure S7C).

Proline 154 mutated in line L14 is a conserved residue in ERH1 plant homologues (*Enhancing RPW8-MEDIATED Hr-like cell death 1)*. *erh1* mutants display constitutive SA accumulation and *PR1* gene expression (Wang et al., 2008). Hence, we measured *PR1* transcripts by RT-qPCR in response to SA in wild-type and L14 plants. Both lines showed similar *PR1* expression in seedlings under 0 and 0.5 mM SA (Supplementary Figure S7B), indicating that ERH1^P154L^ mutation does not seem to be involved in the L14 phenotype.

### Interaction of HAK5 with NPF6.2 and NPF6.3 proteins

The above results support the notion that the phenotype of L14 plants most likely arise from the combination of the loss of AKT1 and the GoF mutation NPF6.2^V210M^. Because HAK5 becomes the main K^+^ uptake pathway in the *akt1* background, we tested whether proteins HAK5 and NPF6.2 interacted. Indeed, the interaction tested positive in a split-ubiquitin assay in yeast (Figure 6A). The strength of HAK5-NPF6.2 interaction was weaker than that of the HAK5-HAK5 homodimer because it could be disrupted by supplemental methionine that decreased the amount of CubPLV-tagged proteins encoded in plasmid pMetYC. The formation of NPF6.2-HAK5 heteromers was confirmed by a BiFC assay in tobacco leaves (Figure 6B). NPF6.2 and HAK5 homodimers were also observed. Notably, HAK5 was also able to interact with the NO3^-^ transceptor protein NPF6.3/NRT1.1, which is a critical protein in NO3^-^ sensing (Figure 6A).

**Figure 6.**
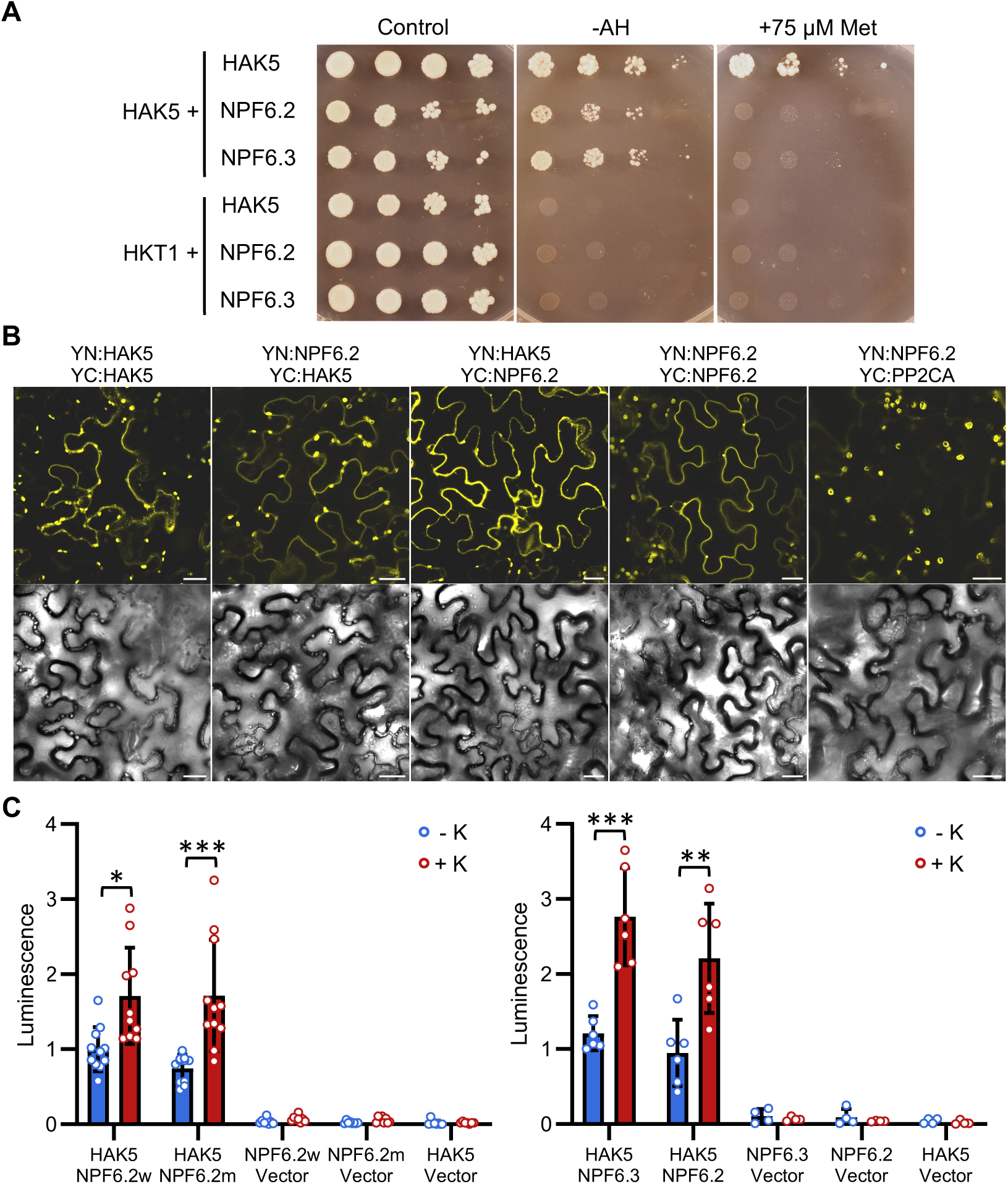
Interaction of HAK5 with NPF6.2 and NPF6.3. **(A)** Split-ubiquitin assay for protein dimerization in yeast. Interaction of NubG- and CubPLV-tagged proteins was detected on selective YNB medium (-AH). Methionine (75 µM) was added to reduce the expression of CubPLV-tagged proteins in plasmid pMetYC to suppress weak interactions. HKT1 was used as negative control. Growth was monitored for 5 days at 28°C. **(B)** BiFC in tobacco. The N-terminus (YN) and C-terminus (YC) of YFP were fused to HAK5 and NPF6.2 as indicated and transiently co-expressed in *Nicotiana benthamiana* leaves. The protein phosphatase PP2CA was used as negative control. Scale bars, 25 µm. **(C)** nLuc reconstitution. Luminescence produced by co-expression of HAK5 and NPF proteins fused to nLucN and nLucC moieties, respectively, in yeast cells in response to KCl. Luminescence reading started after addition of the nanoluciferase substrate and again 15 minutes after supplementation with 50 mM KCl. NPF6.2w and NPF6.2m mean wild-type and V210M mutant proteins. Luminecence values were normalized to that of HAK5-NPF6.2w samples. Shown are the mean values and SE of 6-11 biological replicates. with three technical repeats each. Negative controls were cells co-transformed with an empty vector. Asteriscs indicate statistical differences at *p*<0.05 (*), *p*<0.01 (**), and *p*<0.001 (***) by the *t*-Test.

The reconstitution of nanoluciferase (nLuc) was assayed to quantitate the interaction of HAK5 with the wild-type and mutant NPF6.2 variants, and with NPF6.3. The small size of nLuc (170 aa), and the highly asymmetrical sizes of the complementing fragments (nLuc-N moiety is 159 residues, and nLuc-C is only 11) largely eliminate self-assembly and make this *in vivo* reporter suitable for monitoring protein association in a quantitative manner (Wang et al., 2020). nLuc moieties were fused C-terminally to proteins being tested. Recombinant proteins were expressed in yeast TTE9 cells that are deficient in endogenous K^+^ uptake systems, so that HAK5 was the operative K^+^-uptake system (Ragel et al., 2015). Cells grown in 50 mM KCl were transferred to K^+^-free medium and then re-supplied with KCl. Luminescence was measured before and 15 minutes after KCl addition and normalized to the abundance of the interacting proteins in each condition, determined as the luminescence of HAK5, NPF6.2 and NPF6.3 proteins fused to full-length nLuc (Supplemental Figure S8A). Recordings of reconstituted luminescence showed that HAK5 interacted with both NPF6.2 variants and with NPF6.3, independently of whether nLuc-N and nLuc-C tags were permuted between these proteins (Figure 6C; Supplemental Figure S8B,C). Notably, supplemental K^+^ stimulated these protein interactions. NO3^-^ was not tested because *S. cerevisiae* lacks NO3^-^ uptake proteins and NPF6.2 and NPF6.3 proteins are not functional in this species (Morales de los Ríos et al., 2021). The mutant protein NPF6.2^V210M^ seemed to interact more effectively with HAK5 than the wild-type NPF6.2 when fused to the nLuc-N moiety (*p*=0.02, *t*-test, Supplemental Figure S8B) but not when the nLuc tags were permuted (*p*=0.46, Figure 6C).

Next, we explored whether the interaction of HAK5 and NPF6.2/6.3 proteins modified their transport activity when co-expressed in *Xenopus* oocytes. First, we confirmed protein interaction in oocytes by BiFC; fluorescence was emitted from the oocyte surface indicating protein interaction at the membrane (Supplemental Figure S9). When co-expressed with HAK5, proteins NPF6.2 and NPF6.3 had a minor effect on K^+^ transport measured by TEVC (*p*=0.155 and *p*=0.123, respectively, *t*-test) whereas the mutant protein NPF6.2^V210M^ produced a statistically significant reduction in HAK5-dependent currents (*p*=0.001) (Figure 7A). However, when the transporters were co-expressed with the kinase complex CIPK23/CBL1, the inhibition of HAK5 by NPF6.2 also became significant (*p*=0.001 for NPF6.2; *p*=4.3×10^-4^ for NPF6.2^V210M^), but not for NPF6.3. In the converse experiment, HAK5 completely abrogated the ^15^NO3^-^ uptake mediated by NPF6.2 and NPF6.2^V210M^ (Figure 7B). Together, these results imply that the physical interaction of HAK5 and NPF6.2 results in mutual interference of their transport activity, and that the GoF mutant NPF6.2^V210M^ has a greater inhibitory activity towards HAK5 than the wild-type protein.

**Figure 7.**
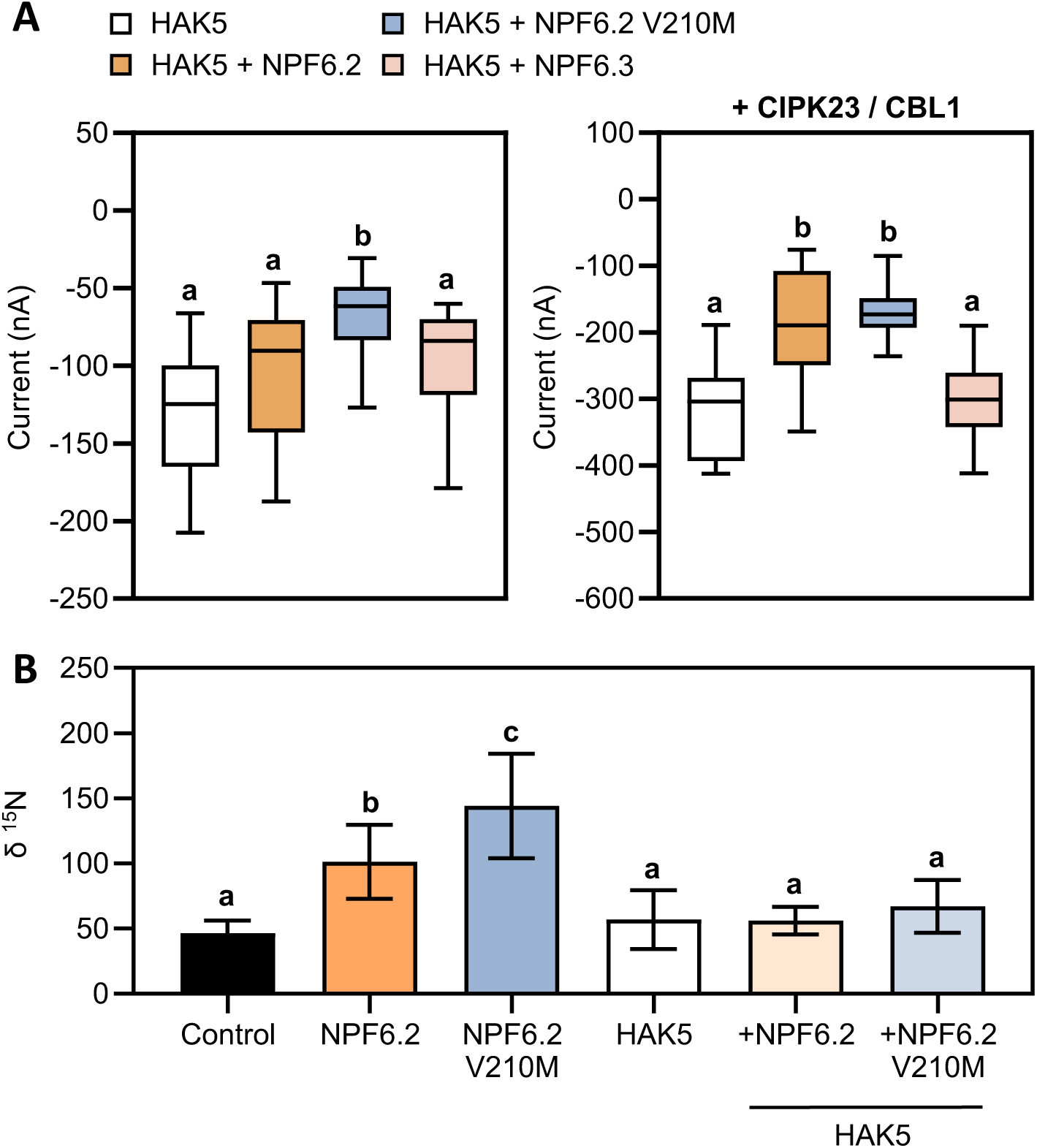
Mutual interference of HAK5 and NPF6.2 proteins in *Xenopus* oocytes. **(A)** NPF6.2 inhibits K^+^ transport by HAK5. TEVC currents after co-expression of HAK5 with NPF6.2, NPF6.2^V210M^ and NPF6.3, with and without the kinase complex CIPK23/CBL1. Mambrane potential was clamped at -120 mV and the bath medium contained 2 mM KCl. Data are peak currents after K^+^ addition (n=10-18). Letters indicate significant differences (*p* < 0.05) by ANOVA comparison. **(B)** HAK5 inhibits nitrate transport by NPF6.2. Water-injected (Control), NPF6.2, NPF6.2^V210M^ and CIPK23/CBL1 with or without HAK5 protein-expressing oocytes were incubated for two hours with 30 mM of ^15^N-labeled KNO3 at pH 5.5. The ^14^N/^15^N ratio was determined in dried oocytes. Data are mean and SD (n=10–15 oocytes). Letters indicate significant differences (*p* < 0.05) by ANOVA with Tukey’s post-hoc test between different batches.

The AlphaFold software (Jumper et al., 2021) allows the accurate prediction of HAK5 and NPF6.2 protein structures, which are highly consistent with the known structures of the bacterial K^+^-H^+^ symporter KimA that is phylogenetically related to HAK5, and the Arabidopsis protein NPF6.3/NRT1.1 that is highly similar to NPF6.2 (Sun et al., 2014; Tascon et al., 2020). Residue V210 of NPF6.2 is predicted to locate at the interaction surface between protomers (Figure 8A). Modeling of a theoretical HAK5-NPF6.2 heterodimer suggested that mutation of residue V210 in NPF6.2 could affect the interaction between these proteins (Figure 8B). Note that the predicted interaction surface of HAK5 with NPF6.2 is on the opposite side to the dimerization domain of HAK5 (Figure 8C). These structural predictions, albeit theoretical, are coherent with the transport results showing that HAK5 abrogated NO3 transport by NPF6.2 and NPF6.2^V210M^, presumably by disrupting the homodimer, whereas NPF6.2 could only partly inhibit the activity of HAK5 because it associates to the opposite side of the HAK5 protomer interaction surface.

**Figure 8:**
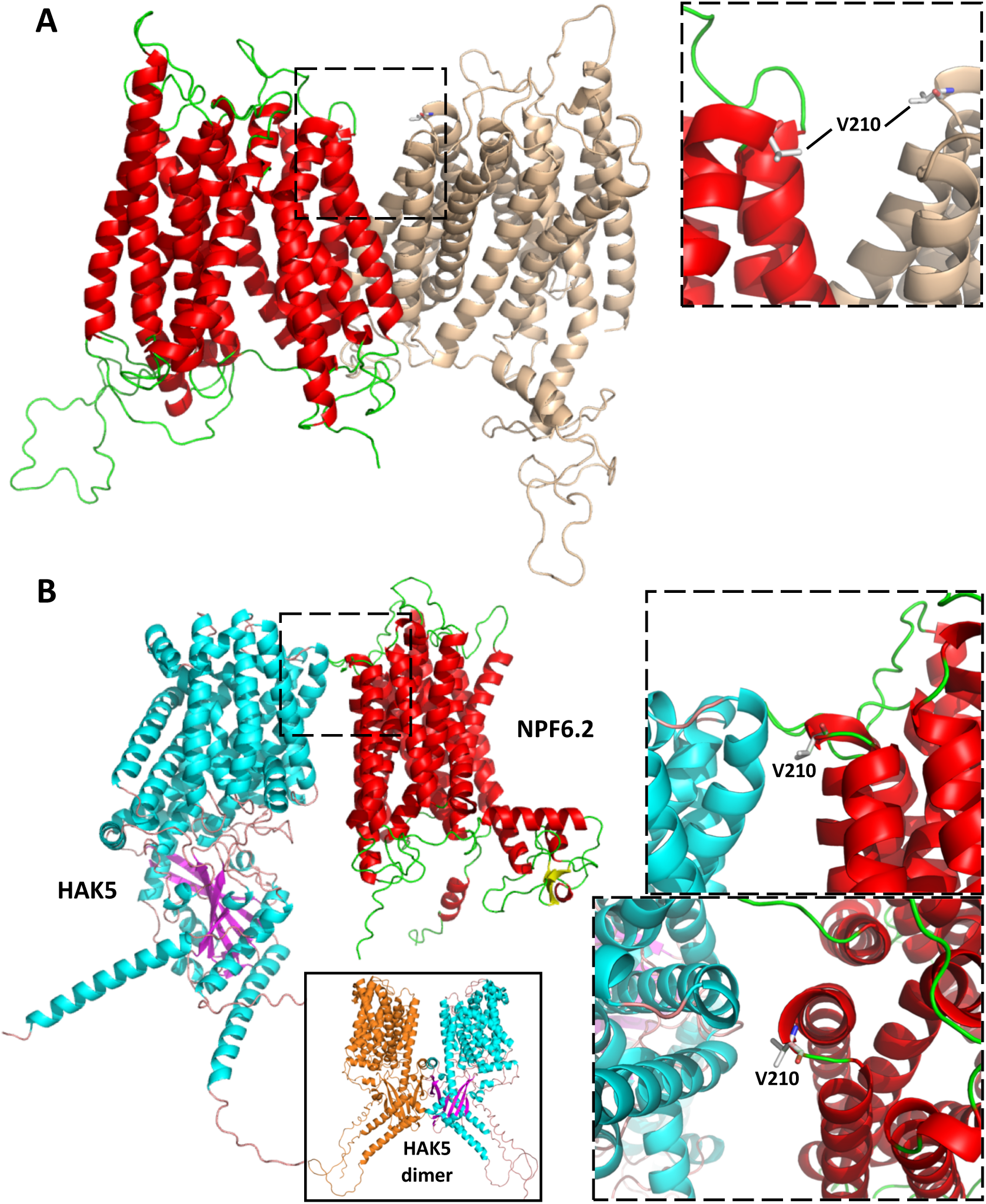
AlphaFold prediction of dimer structures. **(A)** Prediced coupling between NPF6.2 protomers, colored as red/green and bronce. The right panel corresponds to the section marked with the dashed line in the left structure. The position of Val210 in both protomers is marked. **(B)** Left, structure of the putative heterodimer formed between HAK5 and NPF6.2 protomers. The upper rigth box shows the zoomed view of the dashed box. and the lower right panel is the top-view from the cytosolic side. The inset at the bottom center shows the structure of the HAK5 dimer.

## DISCUSSION

A genetic screen aimed to find regulators of *HAK5* gene expression yielded the mutant line L14, which demonstrated stable enhanced expression of the *pHAK5:LUC* reporter under K^+^-sufficient conditions. Mapping-by-sequencing identified two tightly linked mutations in genes involved in K^+^ and NO3^-^ nutrition, namely the loss-of-function (LoF) mutation AKT1^W135Z^ combined with a gain-of-function (GoF) allele NPF6.3^V210M^. AKT1 and HAK5 constitute the two major K^+^-uptake pathways in Arabidopsis roots, as they embody respectively the Low- and High-affinity systems (Nieves-Cordones et al., 2014; Ragel et al., 2015). Loss of AKT1 function is sufficient to induce the expression of *HAK5* owing to the impaired channel-mediated low-affinity K^+^ uptake (Aleman et al., 2011; Ragel et al., 2015). However, comparison of *akt1-2* and L14 plants evidenced that the later had a much stronger phenotype, implying the presence of an additional enhancer mutation. Compared to *akt1-2*, L14 plants had much greatly reduced root K^+^ contents, and were primed for *HAK5* gene induction and high-affinity Rb^+^ uptake without the need of a prior K^+^ starvation step (Figure 4 and Supplemental Figure S4). The enhancer mutation was identified as the GoF allele NFP6.2^V210M^, consistent with the genetic lesion in L14 having a dominant behavior in BC1F1 plants heterozygous for the underlying mutations. Other SNPs fixed in L14 and linked to AKT1^W135Z^ and NPF6.2^V210M^ mutations (Supplemental Table S1) were disregarded based on published data or experimental analyses (Supplemental Figure S8).

The analysis of K^+^ and NO3^-^ contents, and the nutritional status of L14 plants showed that L14 plants had a severe defect in K^+^ uptake and root/shoot partition that could not be explained by the *akt1* mutation alone, and was not matched by a knock-out *npf6.2* mutation (Supplemental Figure S6). These data strongly suggested that the anomalous K^+^ distribution in L14 plants was a consequence of the GoF mutation NPF6.2^V210M^. The analysis of K^+^ and NO3^-^ contents indicated that enhanced NO3^-^ transport by protein NFP6.2^V210M^ acted in synergy with the loss of AKT1 to intensify the nutritional and developmental maladies in L14 plants. Therefore, we propose that the acute phenotype of L14 plants compared to the *akt1-2* mutant results from increased NO3^-^ transport by the mutant protein NFP6.2^V210M^ and with the inhibition of HAK5 transport, which becomes the prevalent K^+^ uptake system in the *akt1* mutant background (Ragel et al., 2015). This conclusion is further supported by the finding that NFP6.2^V210M^ showed enhanced inhibition of HAK5 transport in *Xenopus* oocytes, as discussed below.

Here we show that HAK5, responsible for the high-affinity K^+^ uptake by roots, and the low-affinity nitrate transporter NPF6.2 interact physically and functionally when co-expressed in yeast and *Xenopus* oocytes. NPF6.2 but not NPF6.3 reduced HAK5-dependent K^+^ influx in oocytes, particularly upon the co-expression of the common regulator CIPK23-CBL1. Remarkably, the mutant protein NPF6.2^V210M^ showed a greater interference with HAK5 transport that manifested even in the absence of CIPK23-CBL1. Co-expression of CIPK23-CBL1 minimized the differences between NPF6.2 and NPF6.2^V210M^ in their capacity for HAK5 inhibition. This is coherent with the gain-of-function nature of the NPF6.2^V210M^ mutation. Conversely, HAK5 interfered with nitrate uptake by NPF6.2 and NPF6.2^V210M^ in oocytes. These findings are puzzling considering that K^+^ and N nutrition are tightly and positively linked. Because HAK5 is synthesized under K^+^ limiting conditions whereas NPF6.2 is biochemically active only when NO3^-^ is available, we hypothesize that HAK5 interference with NPF6.2 activity aims to reduce NO3^-^ transport under K^+^-limiting conditions to ensure a commensurate uptake and distribution of these nutrients without further signals involved, thereby enabling a direct, short-term response. This feedback regulation would be absent in K^+^-sufficient conditions where *HAK5* expression is repressed. The converse rationale is less robust, since there is no clear benefit of reducing HAK5 activity through the interaction with NPF6.2 when there is already a low K^+^/ NO3^-^ availability ratio. However, our transport assays indicated that NPF6.2^V210M^ inhibition by HAK5 was almost complete, whereas the inhibition of HAK5 by wild-type NPF6.2 was partial and only in the presence of CIPK23, which becomes activated by either K^+^ or N deficiencies (Figure 8). Hence, HAK5 might not be inhibited *in planta* when NFP6.2 is not targeted by CIPK23 in N-sufficient conditions. Note that N-deficiency is known to induce *HAK5* transcription (Rubio et al., 2014; Meng et al., 2016), and then HAK5 activity could be fine-tuned through the interaction with NO3^-^ transporters.

The physical interactions between K^+^ and NO3^-^ transporters we report here is not without precedents. NPF6.3 interacts with KUP2, and binding results in a conformational change in NPF6.3 with unknown effects in NO3^-^ transport or signaling (Ho and Frommer, 2014). The interaction of TaHAK13 with TaNPF5.10 and TaNPF6.3 proteins of wheat has been reported also, but again the functional consequences of it were not investigated (Run et al., 2022). On the other hand, the Cl^-^/NO3^-^ channels SLAC1 and SLAH3 bind to the inward-rectifying K^+^ channel KAT1 to inhibit K^+^ influx in Arabidopsis guard cells. This serves to impair stomata opening and to achieve efficient closure and evidence the crosstalk at the level of transport proteins (Zhang et al., 2016). Together with these precedents, our findings strongly support the existence of a wide network for physical interactions between dedicated transporters of anionic (NO3^-^, Cl^-^) and K^+^ transporters that could modulate reciprocally their activity to achieve a balanced nutrient acquisition.

A pertinent question is whether HAK5 and NPF6.2 are indeed co-expressed *in planta*. NPF6.2 function was initially linked to the gating of NO3^-^ flux at the petioles and leaf midrib of Arabidopsis seedling (Chiu et al., 2004). The reduction of NO3^-^ contents in the petioles of the *npf6*.2 mutant suggested that NPF6.2 was retrieving NO3^-^ from the xylem entering the leaf lamina (Chiu et al., 2004; Morales de los Ríos et al., 2021). However, single-cell RNA-sequencing has shown that *NPF6.2* is also broadly expressed in cortical and stellar cells (Lhamo and Luan, 2021), implying that NPF6.2 may also contribute to nutrient redistribution. On the other hand, *HAK5* is strongly induced by K^+^-starvation in roots, meaning that both proteins NPF6.2 and HAK5 may be co-expressed in root cells. Theoretically, HAK/KUP and NPF proteins might be subject to clustering within the membrane, which would increase their effective local concentration and increase the probability of interactions. In this regard, NPF6.3:GFP appeared as dispersed diffraction-limited fluorescent spots in the plasma membrane (Zhang et al., 2019). The phosphorylation status of NPF6.3 affected the spatiotemporal dynamics at the plasma membrane. The phosphomimetic form NPF6.3^T101D^ showed fast lateral mobility and membrane partitioning, whereas the non-phosphorylatable NRT1.1^T101A^ showed reduced lateral mobility and protein oligomerization prior to endocytosis via the clathrin-mediated endocytosis and microdomain-mediated endocytosis pathways. In this regard, we have shown that co-expression of CIPK23/CBL1 enhanced the ability of NPF6.2 to interfere with HAK5 transport (Figure 7A).

In conclusion, our data demonstrate the intimate crosstalk between K^+^ and NO3^-^ nutrition by means of the physical interaction between key transporters, and uncover a new layer of mutual regulation in the nutrition of K^+^ and nitrogen. This is coherent with the know induction the *HAK5* gene expression in response to both K^+^ and N starvation (Rubio et al., 2014; Meng et al., 2016).

## MATERIALS AND METHODS

### Plant materials and phenotype analysis

*Arabidopsis thaliana* plants of ecotype Columbia-0 and the reporter line *pHAK5:LUC* (Jung et al., 2009) were used. Knockout mutants in *AKT1* (Rubio et al., 2008) and *NRT1.4 WiscDsLox322H05* (Chiu et al., 2004) have been described. Seeds were stratified for 3 days at 4°C, germinated in 7×7 cm pots containing Jiffy Substrates for the Professional Horticulture and grown in a controlled growth chamber with the day/night regime of 16/8h (long day conditions) or 8/16h (short day conditions), 21/18°C, 60% relative humidity, and 120 μmol m^-2^ s^-1^ PAR. Number of rosette leaves was measured when the basal branch reached 5 cm in length, and number of branches and rosette diameter was calculated 21 days after bolting.

### EMS mutagenesis, mutant screening and sequencing

Approximately 2.5 mg seeds of the *pHAK5:LUC* reporter line in the Col-0 background (Chiu et al., 2004) were treated with 0.3% ethyl methanesulfonate for 8 h at room temperature. After thorough washing with sterile H2O, M1 seeds were sowed in soil. M1 plants were self-pollinated to produce M2 plants, which then were pooled into contingents of 30 individuals, and had their M3 seeds collected. For mutant screening, 3-d-old M3 seedlings were grown in K^+^-sufficient nutrient medium as described by Jung et al. (2009), and the expression of the *pHAK5:LUC* reporter was captured as luminescence. Briefly, seedlings were sprayed with 1 mM luciferin sodium salt (Thermo Fisher Scientific) in 0.01% Triton X-100 solution. After placing the plates in the dark for 5 min, the plates were imaged using a cooled CCD camera system IVIS Lumina II (Caliper Life Sciences). Images were analyzed using Living Image 4.2 software. Plants expressing *pHAK5:LUC* reporter gene in K^+^-sufficient conditions were transferred individually to soil and then self-pollinated to produce M4 seeds. Twenty M4 seeds were grown in K^+^-sufficient medium again, screened by bioluminescent and line L14 was finally selected among M4 plants for further analyses.

Genetic mapping was accomplished with plants collected from a BC1F2 population derived from a cross between line L14 and the parental *pHAK5:LUC*. Twenty individuals with mutant (root luminescence in K^+^-sufficient conditions; see Figure 1) or wild-type phenotypes were pooled separately, genome sequenced, and analyzed by a mapping-by-sequencing approach (James et al., 2013). Genome sequencing was done using the Illumina HiSeq2000 system at 99.8% coverage and 83-92X depth (Macrogen Inc.). After sequence assembly, the reads were aligned to the TAIR10 Arabidopsis Col-0 genome. SNPs enriched in the mutant population relative to the wild-type segregants were identified.

### Arabidopsis nutrient content, ^15^N-nitrate and Rb^+^ uptake assays

Seeds were germinated at room temperature in plastic holders with distilled water and seedlings were transferred to opaque plastic containers for hydroponic culture in controlled growth chambers with the day/night regime of 16/8 (long-day conditions) or 8/16 (short-day conditions) hours, 21/18°C, 60% relative humidity, and 120 μmol m^-2^ s^-1^ PAR. Unless otherwise is stated, plants were grown and treated hydroponically in LAK medium, a modified Long Ashton mineral solution that contains 1 mM K^+^ and 4 mM NO3^-^ (Barragan et al., 2012). The composition of the LAK base solution was: 1 mM KH2PO4, 2 mM Ca(NO3)2, 1 mM MgSO4, 30 µM H3BO3, 10 µM MnSO4, 1 µM ZnSO4, 1 µM CuSO4, 0.03 µM (NH4)6Mo7O24, and 100 µM Fe^2+^ as *Sequestrene* 138-Fe, pH 5.3. When required, supplemental K^+^ was added as K2SO4. For low-K^+^ medium (<1 mM), NaH2PO4 replaced KH2PO4 and K^+^ was added as K2SO4. For experiments in which K^+^ and NO3^-^ concentration were changed simultaneously, 2.5 mM CaCl2 replaced 2 mM Ca(NO3)2 and KNO3 was added as required.

For Rb^+^ uptake experiments, plants grown in LAK medium with 1 mM K^+^ were transferred to a LAK without K^+^ (NaH2PO4 replaced KH2PO4) for 72 hours. Individual plants were then transferred to nutrient solution with no K^+^ added and supplemented with 50 µM RbCl. After 2 h, plants were harvested, roots and shoots were separated, and dried at 65°C for 3 days.

K^+^ and Rb^+^ concentrations were determined in acid-extracted samples (nitric acid 65% [v/v], hydrogen peroxide 30% [v/v]) by Atomic Emission Spectrometry. For determination of nitrate content, about 3 mg of dry tissue per sample were mixed with 1 ml of miliQ water and incubated at 60°C for 30 minutes. NO^-^3 present in the sample were measured by High Pressure Ion Chromatography (HPIC). For ^15^N-nitrate uptake assay, plants were preincubated for 2 min in 0.1 mM CaSO4 and transferred for 5 min in LAK base solution containing 0.5 or 5 mM K^15^NO3 (99.9% ^15^N abundance, Sigma-Aldrich). Roots were then washed in 0.1 mM CaSO4, dried at 65°C and ^15^N content was analyzed by Continuous Flow Mass Spectrometry, using a VarioPyroCube elemental analyzer (Elementar, UK) coupled with an IsoPrime Precision mass spectrometer (Elementar, UK).

### Quantitative Real-Time RT-PCR Analysis

Plants were grown in hydroponic culture in short-day conditions. After 5 weeks, plants were harvested and total RNA was isolated using ISOLATE II RNA Plant Kit (Bioline). Two micrograms of DNA-free RNA was then reverse-transcribed using the QuantiTect® Reverse Transcription Kit (QIAGEN). Quantitative real-time RT-PCR analysis was per-formed using the iCycler iQ™5 system and iQ SYBR® Green Supermix (Bio-Rad Laboratories). All primers used in this work are listed in Supplemental Table S2. Primers HAK5-F and HAK5-R were used to amplify the endogenous *HAK5* transcripts. For the normalization of *HAK5* transcripts, *Ubiquitin 10* primers (UBQ10-F and -R) were used as an internal control. The fold change of *HAK5* transcript in each sample was calculated based on the efficiency calibrated model (Yuan et al., 2006) and compared with the *HAK5* transcript level of the wild-type control under the K^+^-sufficient condition.

### Yeast methods

The transport activity of AKT1 and AKT1^W135Z^ was tested in the yeast *Saccharomyces cerevisiae*. The AKT1^W135Z^ coding region was amplified from L14 mutant line by PCR using primers AKT1-F and -R. The amplicon was cloned between *XmaI* and *EcoRI* in the p414GPD expression vector (Mumberg et al., 1995). *AKT1* (*At2g26650*) from pFL61-AKT1 (Sentenac et al., 1992) was cloned in vector pRS415 using *XbaI* and *BamHI* restriction sites. Plasmids were transformed into the yeast strain TTE9.3 (*MATa, ena1Δ::HIS3::ena4Δ, leu2, ura3-1, trp1-1, ade2-1, trk1Δ, trk2::pCK64*) (Bañuelos et al., 1995; Bañuelos et al., 2002) using the PEG-lithium acetate protocol (Gietz and Schiestl, 2007). For growth complementation assays, 5 µL aliquots from yeast cultures (DO600 = 1) or 10-fold serial dilutions were spotted onto AP plates (Rodriguez-Navarro and Ramos, 1984) supplemented with 1 or 50 mM KCl. Plates were incubated at 28°C for 3-4 days.

For expression in *Hansenula polymorpha*, the *NPF6.2^V210M^* mutation was performed in pYNR-EX-NPF6.2 (Morales de los Ríos et al., 2021) using Q5® Site-Directed Mutagenesis Kit (New England Biolabs). Strain *Δynt1*, a derivative of NCYC495 (*leu2 ura3*), was used for genetic complementation with NPF6.2 and NPF6.2^V210M^ (Martín et al., 2008). Plasmids were transformed by electroporation as described (Morales de los Ríos et al., 2021). Single yeast colonies were grown 16h at 37°C and shaking in YGNH medium (Yeast Nitrogen Base 0.17% (w/v), 2% glucose, 5 mM NH4Cl as nitrogen source) (Martín et al., 2008). For cell growth experiments, centesimal and millesimal dilutions of cell cultures at DO660 ∼ 1 were incubated in the same medium without nitrogen source (YG) plus 0.5 mM KNO3 for 48 h. Yeast growth was measured in a microplate-reader Varioskan™ LUX (Thermo Scientific) and expressed as OD660.

### Split-ubiquitin

The *S. cerevisiae* strain THY.AP4 (*MATa ura3 leu2 lexA::lacZ::trp1 lexA::HIS3 lexA::ADE2*) and the yeast shuttle vectors pNXgate32-3HA and pMetYCgate have been described (Obrdlik et al., 2004). The Arabidopsis *HAK5, NPF6.2 and NPF6.3* open reading frames (ORFs) without the stop codon were amplified with Accuzyme high-fidelity DNA polymerase (BIOLINE) with primers attB1_HAK5-F and attB2_HAK5-R for *HAK5,* attB1_NRT1.4-ATG and attB2_NRT1.4-stop for *NPF6.2* and attB1_NRT1.1-ATG and attB2_NRT1.1-stop for *NPF6.3.* For NubG fusions, the pNXgate32-3HA vector was cleaved with *Eco*RI/*Sma*I and used together with PCR products to transform yeast cells for *in vivo* cloning (Fusco et al., 1999). Transformants were selected on SD minimal medium lacking tryptophan (T). For CubPLV fusions, the vector pMetYCgate was cleaved with *Pst*I/*Hind*III and used together with PCR products to transform yeast. Transformants were selected on SD lacking leucine (L). Several yeast clones from each transformation were selected and plasmids were rescued in *E. coli* TOP10 and purified to verify by sequencing the correct cloning of cDNAs. For the interaction assays, plasmids pNXgate32-HA and pMetYCgate containing cDNAs were co-transformed in the yeast strain THY.AP4 and transformants were selected on YNB-LT. Over-night cultures of the yeast transformants were grown in liquid YNB−LT medium and 10-fold serial dilutions were spotted on YNB plates lacking leucine, tryptophan, adenine and histidine (YNB−LTAH). Plates were incubated at 30°C for 4 d. The full length HKT1 cloned in pMetYC vector was used as control of interaction.

### NanoLuciferase reconstitution

We developed vectors for split-nanoLuciferase complementation in yeast. To allow *in vivo* cloning of the gene of interest, the attB1-KanMX-attB2 cassette of pNXgate32-HA (Obrdlik et al., 2004) was amplified by PCR with primers attB1-F and attB2GSG-R. The DNA encoding the N-terminal region of nanoluciferase (nLucN) was amplified by PCR with primers attB2GSG-F and HA2x-R using the vector HBT-nLucN-HA (Deng et al., 2020) as template. The coding sequence of the C-terminal region of nanoluciferase (nlucC) was amplified with primers attB2GSG-F and FLAG2x-R using the vector HBT-nLucC-FLAG (Wang et al., 2020) as template. The PCR fragment containing the attB1-KanMX-attB2 cassette was fused to the nLucN by overlapping PCR using the primers attB1-F and HA2x-R. The final PCR product was digested with *Xho*I/*Aat*II and subcloned in the yeast shuttle vector pDEST32 (Invitrogen) cleaved with the same restriction enzymes; the resulting construct was named p32nLucNgate. In the same way, the attB1-KanMX-attB2 cassette was fused to the nLucN with primers attB1-F and FLAG2x_rev and subcloned in pDEST22 (Invitrogen) producing the plasmid p22nLucCgate. For protein fusions with full-length nLuc the p22nLuc plasmid was designed. nLuc moieties were PCR amplified from p32nLucN HAK5 with the oligos HAK5-830 and nLuc-P2, and from p22nLucC gate with the oligos M13-F and nLuc-P1, and assembled by fusion PCR with oligos HAK5-830 and M13-F. The PCR product, digested with BglII-SacI, was cloned into the corresponding sites of p22nLucC-HAK5. Next, Arabidopsis *HAK5, NPF6.2 and NPF6.3* ORFs without the stop codon were amplified as described above. The purified PCR products were cloned *in vivo* (Fusco et al., 1999) in vectors p22nLuc, p32nLucNgate and p22nLucCgate by co-transformation of the yeast strain TTE9 (*trk1, trk2*) or the wild-type W303 (*TRK1 TRK2*).

For the split-nanoLuciferase complementation assays, yeast transformants expressing proteins tagged with nLuc moieties at their C-termini were grown in AP minimal medium supplemented with 50 mM KCl. Cells were incubated in AP without KCl 2-4 hours, after which 90 µl samples were mixed with 10 µl of Nano-Glo® Luciferase assay reagent (Promega, Madison WI) and luminescence was measured every 30 seconds for 5 minutes to establish the baseline. Cells were then supplied with 50 mM KCl and luminescence was recorded after 15 minutes, using a Varioskan LUX Multimode Microplate Reader (Thermo Scientific). As controls for nLuc assays, yeast cells were transformed with proteins fused to the full-length nLuc and processed as described for split-nanoLuciferase complementation assays.

### Bimolecular Fluorescence Complementation (BIFC) assays

The cDNAs of *NPF6.2* and *HAK5* were cloned in Gateway vectors (Gehl et al., 2009), generating the constructs pVYNE(R)-NPF6.2, pVYCE(R)-NPF6.2 and pVYCE(R)-HAK5. Plasmid SPYCE(M) containing the PP2CA cDNA has been described (Waadt et al., 2019). Proteins were transiently expressed in *N. benthamiana* leaf epidermal cells through *Agrobacterium tumefaciens* infiltration. Three days after infiltration, YFP fluorescence signals were detected at an excitation wavelength of 515 nm using a confocal laser scanning microscope (Olympus FluoView FV1000).

For BiFC of YFP-tagged proteins in *Xenopus* oocytes, the cDNAs encoding HAK5, NPF6.2 and NPF6.2^V210M^ were cloned into expression vectors based on pNB1u (Nour-Eldin et al., 2006) adapted for BiFC, by the USER (Uracil-Specific Excision Reagent) cloning technique (Jørgensen et al., 2017). Primers used for cloning HAK5 with YFP fusions were HAK5-YFP-F and HAK5-YFP-R. Primers used for cloning NPF6.2 and NPF6.2^V210M^ with YFP fusions were NPF6.2-YFP-F and NPF6.2-YFP-R. Complementary RNA (cRNA) was prepared with the AmpliCap-Max™ T7 High Yield Message Maker Kit (CellScript). For BiFC, 20 ng of YC:HAK5 (vector YCpNB1u) and of YN:NPF6.2 or YN:NPF6.2^V210M^ (vector YNpNB1u) cRNA were injected. The *NPF6.2^V210M^* mutation was performed in pNB1u-NPF6.2 (Morales de los Ríos et al., 2021) using Q5® Site-Directed Mutagenesis Kit (New England Biolabs).

### 15N-nitrate accumulation assay in Xenopus oocytes

For ^15^N uptake experiments, oocytes were obtained and cRNA- or water-injected as previously described (Lacombe and Thibaud, 1998). Oocytes were maintained at 18 °C in ND96 solution (96 mM of NaCl, 2 mM of KCl, 1 mM of CaCl2, 1 mM of MgCl2, and 10 mM of HEPES/Tris at pH 7.4) until use 1-2 days later. For the ^15^N-nitrate assay, oocytes were incubated 2 h in 2 mL of ND96 medium pH 5.5 (5 mM MES) containing between 1 and 30 mM of K^15^NO3 to test influx activity (Léran et al., 2015). For pH-dependency experiments, ND96 solution was adjusted at pH 5.5 (MES), 6.5 (MES-HEPES) or 7.5 (HEPES). For high-K^+^ experiments, ND96 medium pH 5.5 was modified to contain 50 mM KCl or 50 mM NaCl. After incubation, oocytes were washed 5 times in 15 mL of ND96 medium (pH 5.5) at 4°C and dried at 65°C during 24 h. Oocytes were then analyzed for ^15^N abundance by Continuous Flow Mass Spectrometry as described above.

### Electrophysiology in Xenopus oocytes

The cDNAs of interest were inserted downstream of the T7 promoter in the vector pNB1u (Nour-Eldin et al., 2006) by the USER™ cloning method. Constructs pNB1u-AKT1, pGEMKN-CIPK23 and pGEMKN-CBL1 have been described elsewhere (Geiger et al., 2009; Held et al., 2011; Sanchez-Barrena et al., 2020). Plasmids were linearized, and transcribed in vitro with HiScribe T7 ARCA mRNA Kit (New England Biolabs).

Stage V and VI oocytes purchased from Ecocyte Bioscience Europe were injected with 20 ng of AKT1, AKT1^W135Z^, HAK5, NPF6.3, NPF6.2, and NPF6.2^V210M^, along with 4 ng each of CIPK23 and CBL1. After injection, oocytes were incubated for 1–3 days at 16 °C in ND96 pH 7.4 solution. Electrophysiological recordings were performed 1-2 days post-injection using the Two-Electrode Voltage-Clamp (TEVC) technique at a holding potential of -120 mV (for HAK5) or -180 mV (for AKT1). The standard bath solution consisted of 10 mM citric acid/Tris (pH 4.0), 1 mM CaCl2, 1 mM MgCl2, and 1 mM LaCl3. To assess AKT1 and HAK5 transport activity, 2 mM KCl was added to the standard bath solution. The osmolarity of all solutions was adjusted to 220 mOsm/kg with D-sorbitol.

## ACCESSION NUMBERS

The accession numbers were all obtained from the Arabidopsis Information Resource (TAIR). AKT1 (At2g26650), HAK5 (At4g13420), NPF6.2 (At2g26690), NPF6.3 (At1g12110), CIPK23 (At5g47100), CBL1 (At4g17615).

## SUPPLEMENTAL DATA

**Supplemental Figure S1.**
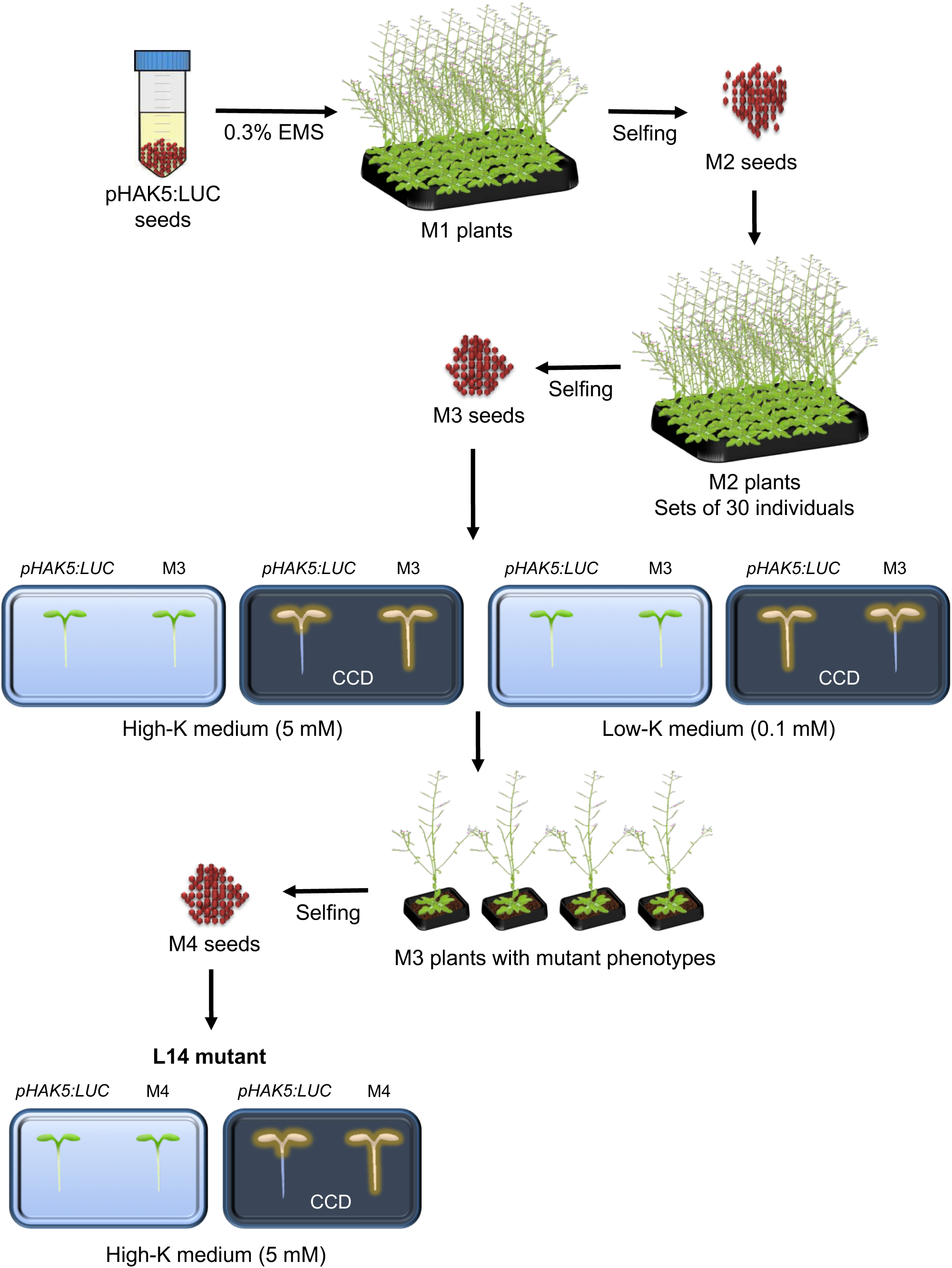
Screening of EMS-mutagenized *pHAK5:LUC* by altered luminescence. An EMS-mutagenized population of the Arabidopsis line expressing the reporter gene *pHAK5:LUC* was screened for altered expression of luciferase under low-K^+^ (0.1 mM) and high-K^+^ (5 mM) conditions. Mutant L14 was selected for further analysis.

**Supplemental Figure S2.**
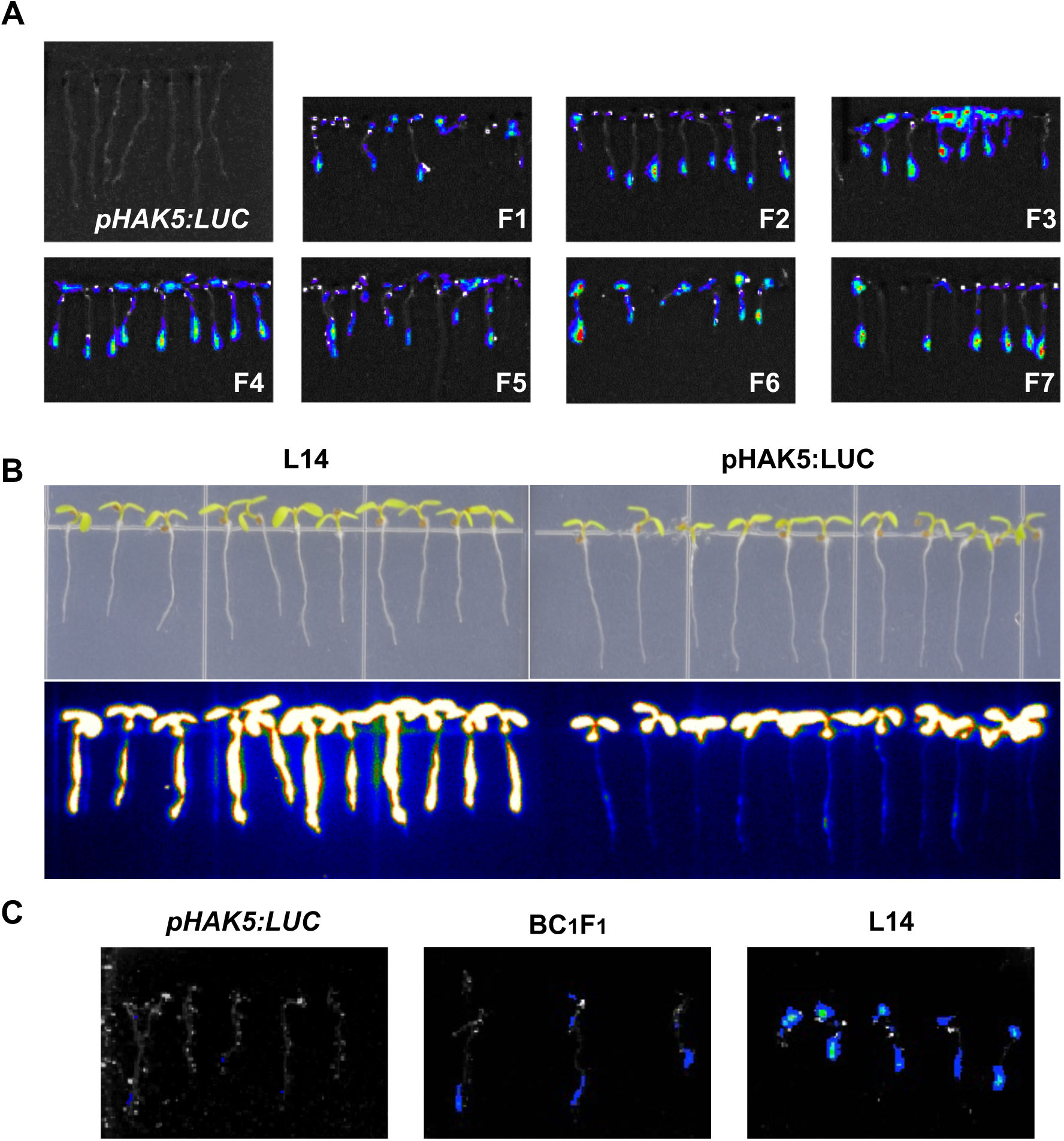
*pHAK5:LUC* expression in wild-type, L14-1 and BC1F1 seedlings. **(A)** 5-day old L14-1 seedlings from generations F1 to F7 were grown in LAK medium with 1 mM KCl. Stronger luminescence signals were detected in L14 mutant seedlings compared to the parental line with *pHAK5:LUC*. **(B)** Luminescence of parental and mutant seedlings in ½ MS medium. **(C)** Expression of the *pHAK5:LUC* reporter gene in seedlings of *pHAK5:LUC*. L14 and BC1F1 lines. Plants were grown on LAK 4 mM K^+^ medium. The expression of the reporter gene in F1 individuals from the cross of line L14 (F5 generation) with its parental *pHAK5:LUC* was analyzed, as well as the expression in the parents. The signal was quantified in an equal surface area (ROI) in the root apex of analyzed individuals, obtaining radiance values (p/s/cm^2^/sr) of 3.2±0.7 (x10^6^) for line L14 (n=7). 2.2±0.6 (x10^6^) for line BC1F1 (n=7) and 3.4±0.3 (x10^4^) for line *pHAK5:LUC* (n=3).

**Supplemental Figure S3.**
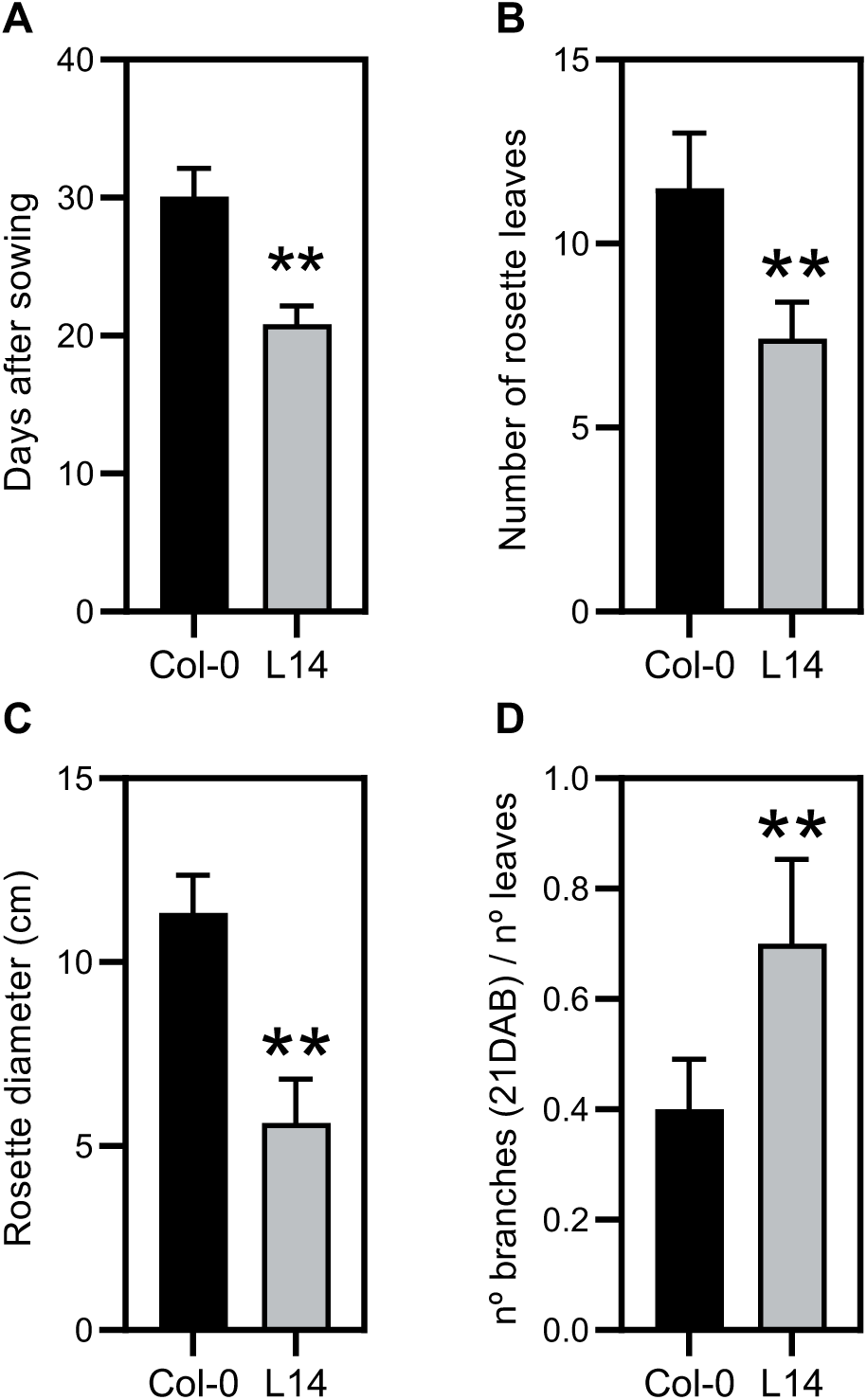
Developmental deffects of L14 plants. WT and L14 plants grown in a controlled growth chamber with the day/night regime of 16/8 hours, 23/20 °C, 70% relative humidity, and 120–140 μmol m-2 s-1 PAR. **(A)** flowering time, **(B)** number of rosette leaves at bolting, **(C)** rosette diameter at 21 DAB and **(D)** ratio of number of branches/leaves. Data are mean and SD (n = 10–12). **p < 0.01 by t-test.

**Supplemental Figure S4.**
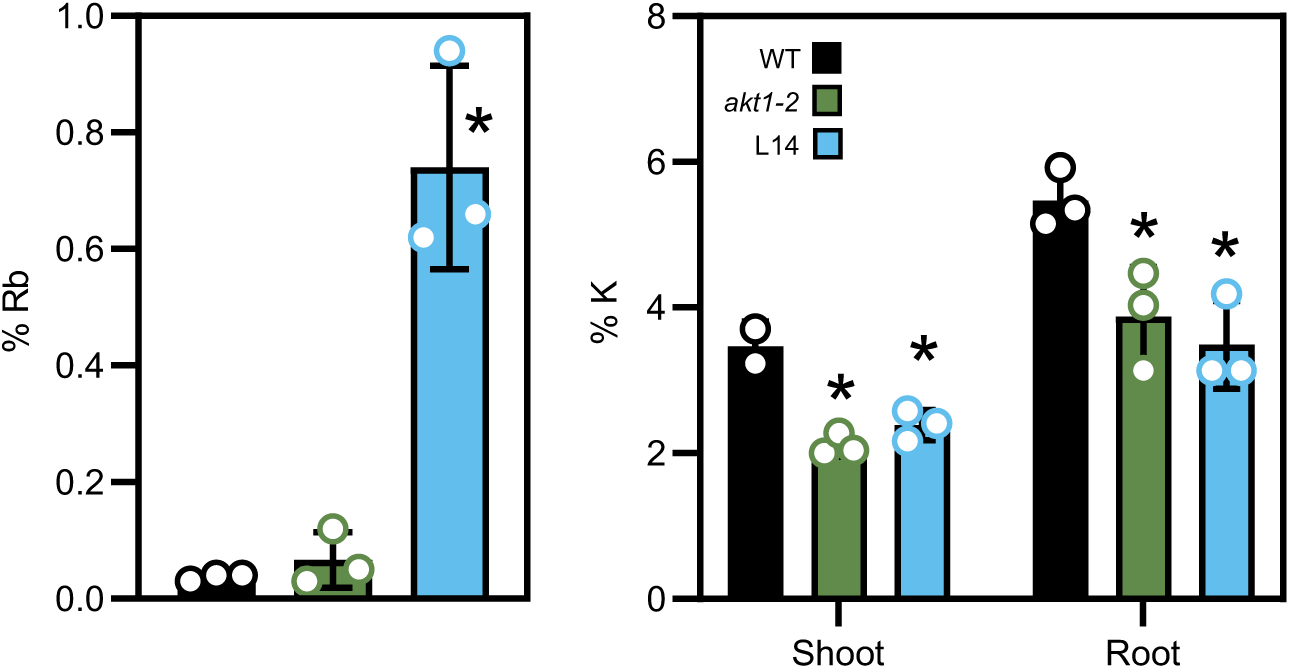
(new). Rb^+^ and K^+^ content in WT, *akt1-2* and L14 lines. Rb^+^ and K^+^ content in WT, *akt1-2* and L14 lines. Plants were grown for 30 days on LAK medium (1 mM KCl). After washing the roots with water, plants were transferred to LAK medium without KCl and supplemented with 50 µM RbCl. After 2 hours, plants were harvested, the roots were washed with cold water and the Rb^+^ **(B)** and K^+^ **(C)** contents in dry matter were determined. Mean and standard deviation values are shown (n= 3 plants). Statistical differences of mutant lines relative to the WT were determined by ANOVA (Tukey’s test),* means *p*<0.05.

**Supplemental Figure S5.**
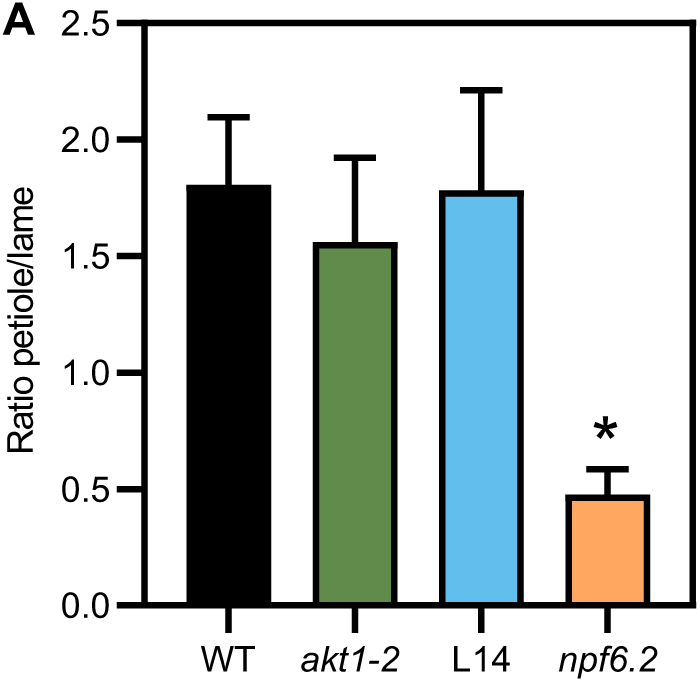
NPF6.2 affects nitrate content in petioles. Ration of nitrate content in petiole and lamina of 3-week-old plants grown in substrate soil. The histograms show the mean +/- SD (n=3). Statistical differences with the WT were determined by ANOVA with Tukey’s post-hoc test (n=3), * means *p*<0.05

**Supplemental Figure S6.**
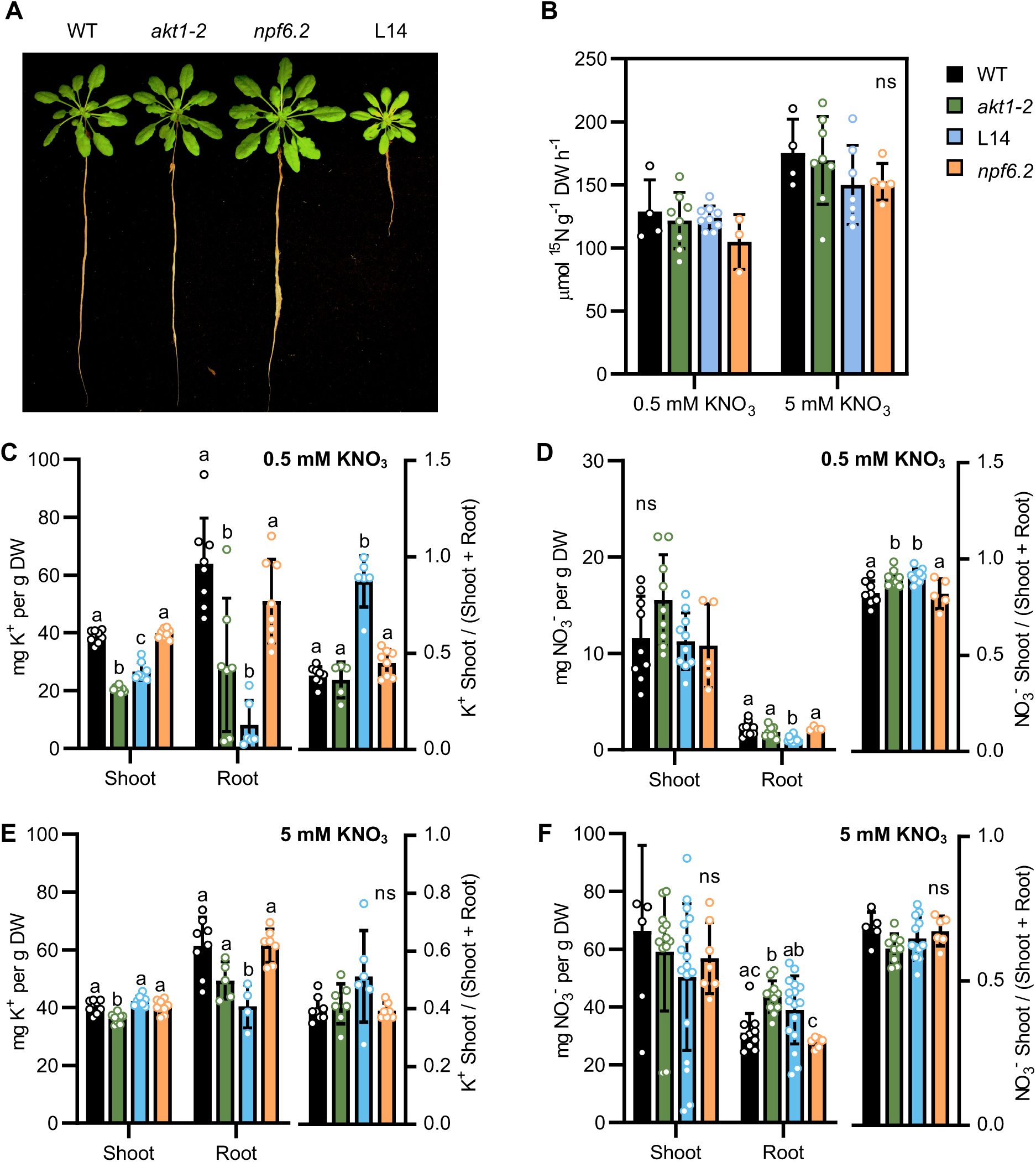
(NEW). Potassium and nitrate content, and ^15^N uptake. **(A)** Plants of the indicated gentotypes grown in LAK medium with 1 mM KNO_3_. **(B)** After 3 weeks in 1 mM KNO_3_. plants were grown for 5 days in fresh medium with 0.5 mM or 5 mM, and trasferred to the same medium with 0.5 mM or 5 mM of ^15^N-labelled KNO_3_ for 5 minutes. The amount of ^15^N taken up was determined in roots. **(C-F)** Plants grown in LAK medium supplemented with 1 mM KNO_3_ for 3 weeks were tranferred to 0.5 mM **(C,D)** or 5 mM **(E,F)** KNO_3_ for 5 days. Bars are mean and SD (n=6-11). Letters indicate significant differences between genotypes at *p*< 0.05 by ANOVA with Tukey’s or Welch’s post-hoc test.

**Supplemental Figure S7.**
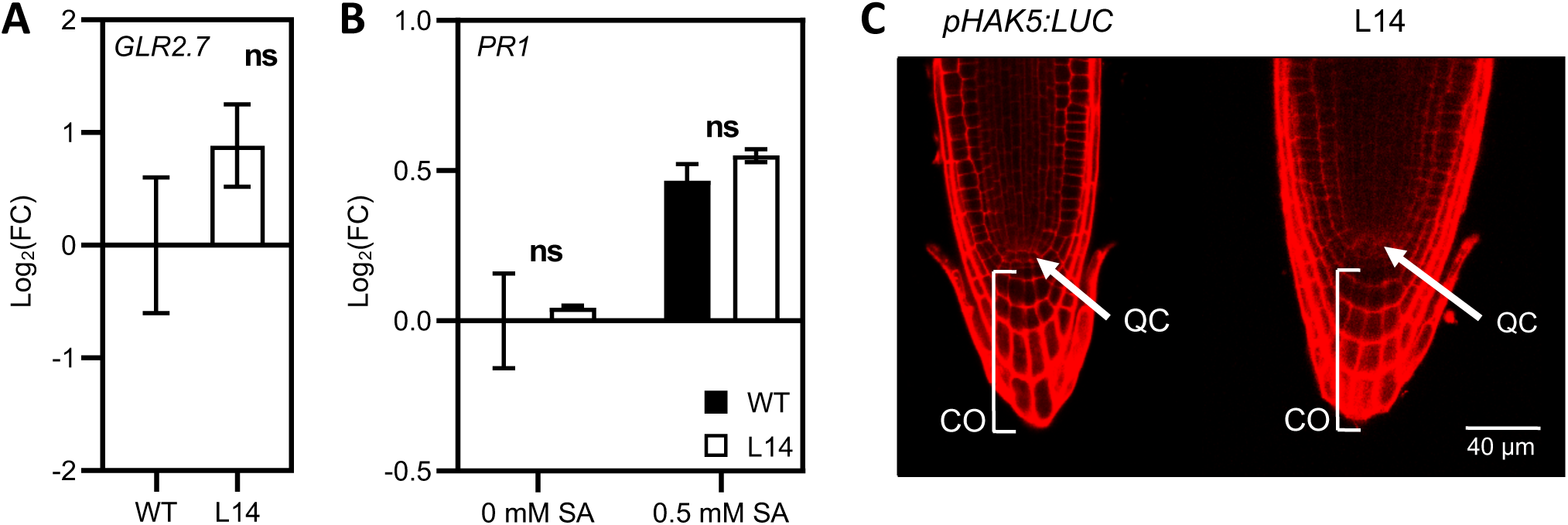
Mutations in GLR2.7, ERH1 and CDF4 do not influence the L14 phenotype. **(A)** Expression of the endogenous *GLR2.7* gene in *pHAK5:LUC* and L14 lines. Expression was quantified in shoots of adult plants. *UBQ10* was used as reference gene and *GLR2.7* expression was normalized to pHAK5:LUC. Differences between lines were not statistically significant (*t-*Test). **(B)** Expression of the endogenous *PR1* gene in *pHAK5:LUC* and L14 lines. Expression was quantified in 7-days plants growth in ½MS. *UBQ10* was used as reference gene and *PR1* expression was normalized *to pHAK5:LUC* (0 SA). Differences in *PR1* expression between lines were not significant (*t-*Test). **(C)** The root tips of 5-day-old seedlings grown in ½MS medium were stained with propidium iodine. The images were taken with a confocal microscope, with an excitation wavelength of 540 nm and emission wavelength of 600-650 nm. Quiescent center (QC), columella (CO)

**Supplemental Figure S8.**
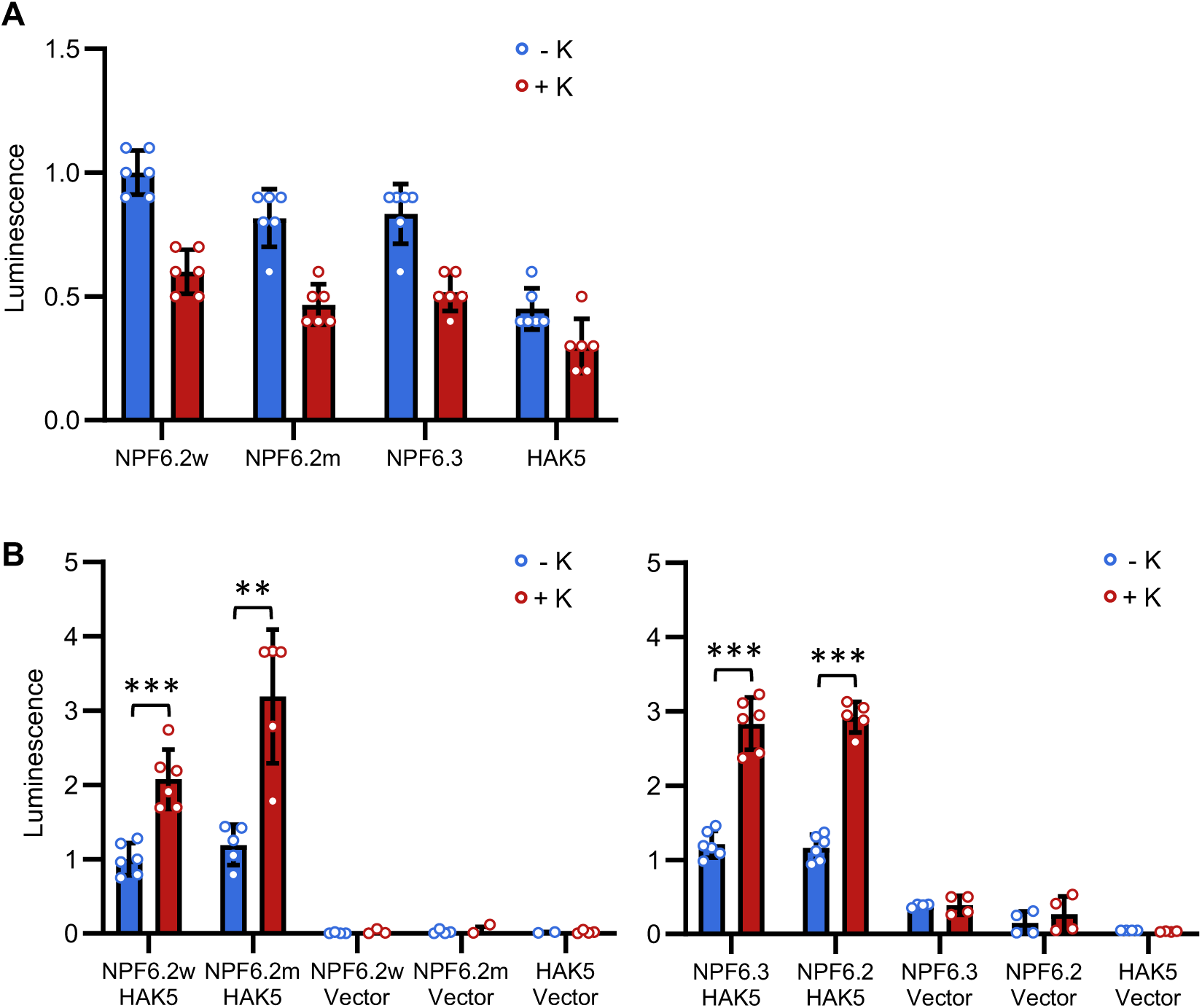
Interaction of HAK5 with NPF6.2 and NPF6.3. **(A)** Lumininescence produced by proteins HAK5, NPF6.2 (NPF6.2w), mutant NPF6.2^V210M^ (NPF6.2m) and NPF6.3, all fused to full-length nLuc, before and after being treated with 50 mM KCl for 15 minutes. Luminescence was normalized to NPF6.2w readings. Shown are the mean values and SE of six biological replicates, with three technical repeats each. **(B)** nLuc reconstitution. Luminescence produced by co-expression of NPF and HAK5 proteins fused to nLucN and nLucC moieties, respectively, in yeast cells in response to KCl. Luminescence reading started after addition of the nanoluciferase substrate and again 15 minutes after supplementation with 50 mM KCl. NPF6.2w and NPF6.2m mean wild-type and V210M mutant proteins. Luminecence values were normalized to that of NPF6.2w-HAK5 samples. Shown are the mean values and SE of six biological replicates, with three technical repeats each. Negative controls were cells co-transformed with an empty vector. The experiment was repeated twice with similar results. Asteriscs indicate statistical differences at *p*<0.01 (**) and *p*<0.001 (***) by the *t*-Test.

**Supplemental Figure S9.**
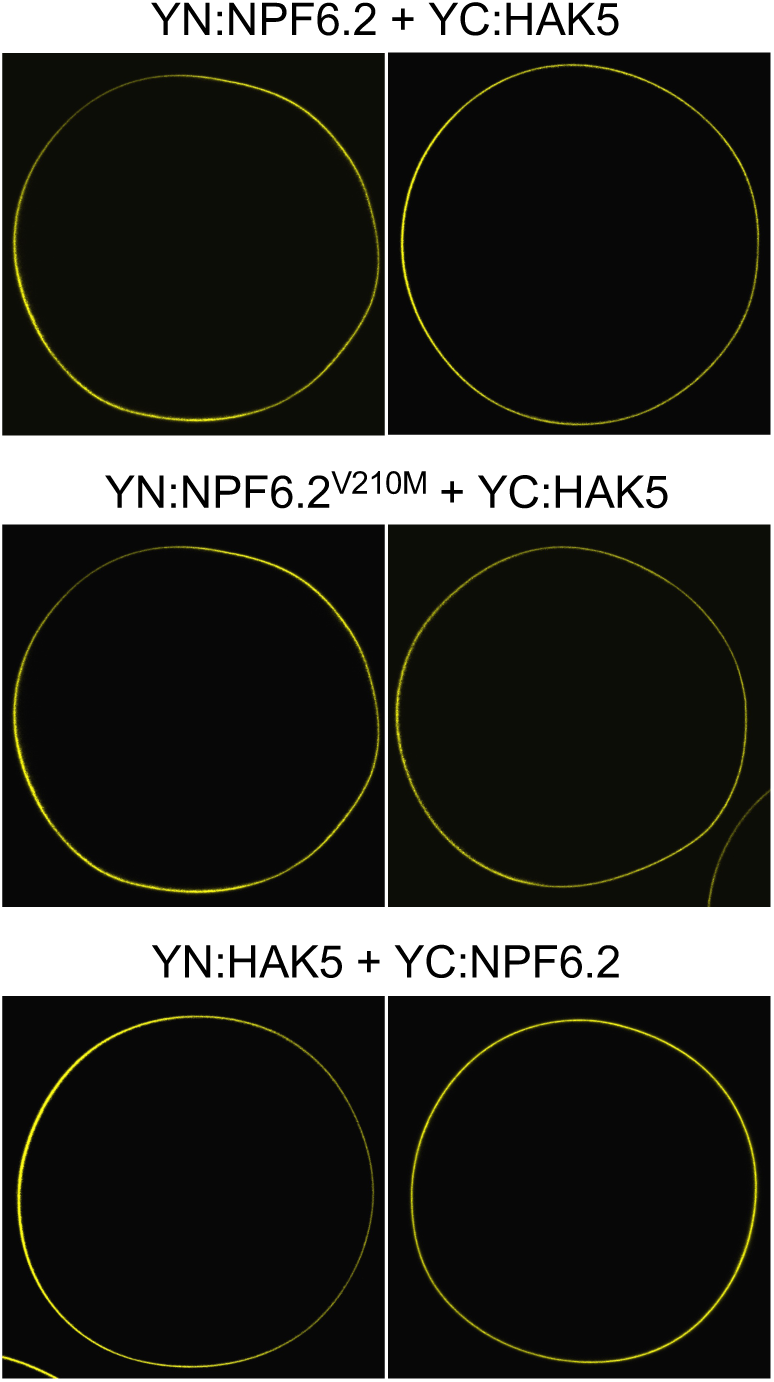
BiFC of HAK5 and NPF6.2 proteins in *Xenopus* oocytes. N-terminal (YN) and C-terminal (YC) moieties of YFP were fused translationally to the N-termini of HAK5, NPF6.2 and NPF6.2^V210M^ proteins in vector pNB1u and expressed in oocytes. Fluorescence was observed by confocal microscopy 2 (left panels) and 3 days (right panels) after cRNAs co-injection. Water-injected oocytes had no fluorescence.

**Supplemental Table S1.** Mutations found in chromosome 2 of line L14. Shown is the genomic position of the base change, the amino acid change involved, the mutant and wild-type allele frequencies (SNPs) in the L14 mapping population, the AGI codes of genes affected, and the proteins encoded.

| Genomic position | Amino acid change | In mutant population |  | Gene | Protein |
| --- | --- | --- | --- | --- | --- |
|  |  | Frequency of mutant allele (%) | Frequency of WT allele (%) |  |  |
| 2590275 | S/L | 77.2 | 42.5 | AT2G06530 | VPS2.1 |
| 6407909 | E/Q | 87.2 | 39.8 | AT2G14910 | Unknown |
| 6997367 | G/E | 75.8 | 57.4 | AT2G16120 | PMT1 |
| 7918254 | D/N | 85.2 | 54.3 | AT2G18193 | Unknown |
| 8313166 | P/S | 88.06 | 0 | AT2G19160 | Transferase |
| 8350250 | G/R | 87.14 | 0 | AT2G19240 | Unknown |
| 8442256 | S/F | 92.5 | 48.57 | AT2G19490 | RECA2 |
| 8641087 | R/H | 98.55 | 41.94 | AT2G20010 | Unknown |
| 10132001 | G/E | 100.0 | 0 | AT2G23800 | GGPPS4 |
| 10541983 | - | 100.0 | 45.45 | AT2G24755 | Unknown |
| 10784032 | G/S | 100.0 | 39.7 | AT2G25320 | TRAF-like |
| 11334975 | W/Z | 100.0 | 39.2 | AT2G26650 | AKT1 |
| 11348616 | V/M | 100.0 | 37.5 | AT2G26690 | NPF6.2 |
| 11975562 | P/L | 100.0 | 39.0 | AT2G28100 | FUC1 |
| 12515358 | - | 100.0 | 41.7 | AT2G29120 | GLR2.7 |
| 12604568 | S/F | 100.0 | 56.5 | AT2G29360 | NAD(P)-binding |
| 13357371 | R/H | 98.25 | 42.31 | AT2G31320 | PARP1 |
| 14414283 | E/K | 87.1 | 0 | AT2G34140 | CDF4 |
| 15743149 | G/D | 85.51 | 52.94 | AT2G37510 | RNA-binding |
| 15878483 | P/L | 86.4 | 41.9 | AT2G37940 | ERH1 |

**Supplemental Table S2.**
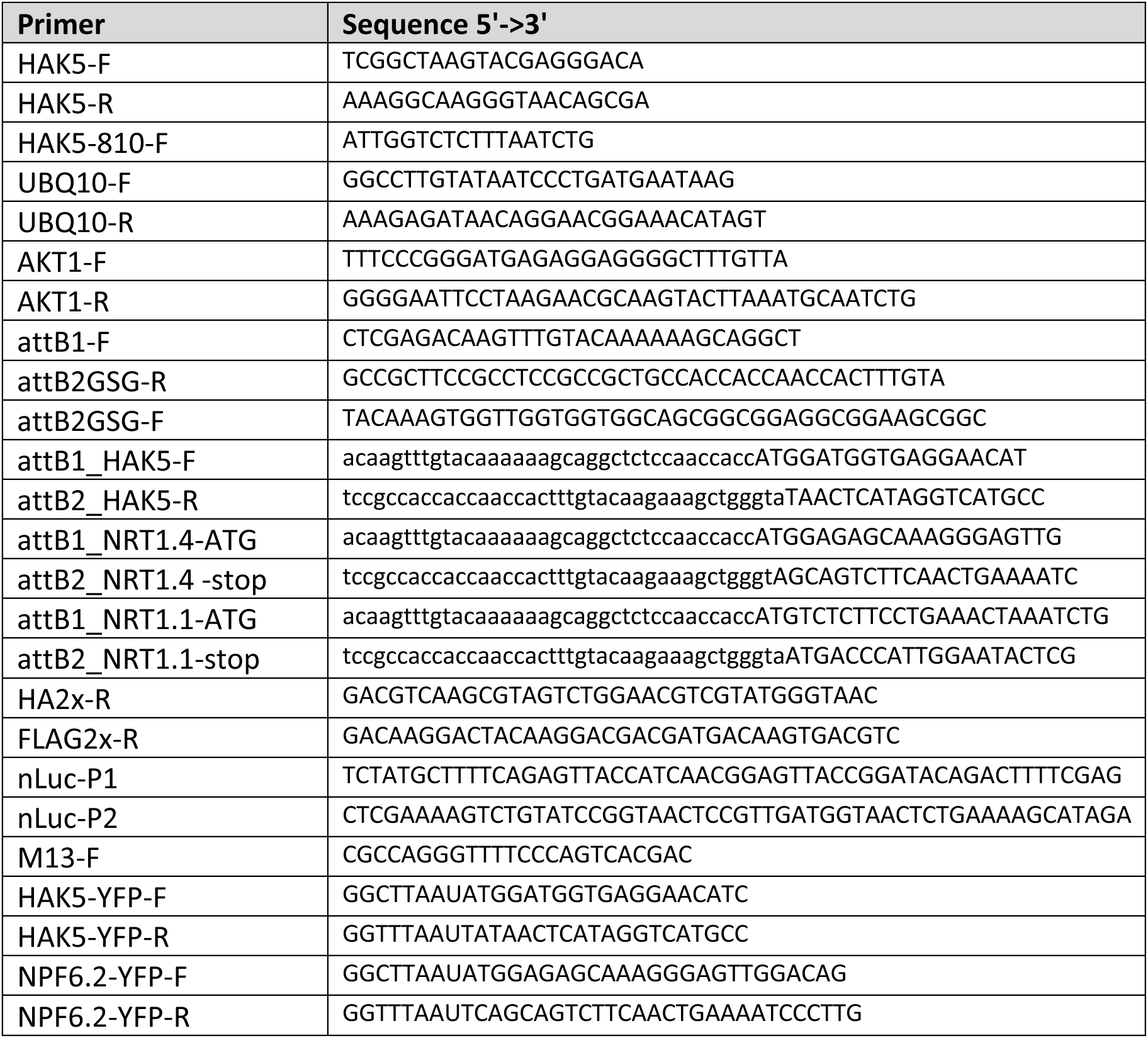
Primers used in this study.

## AUTHOR CONTRIBUTIONS

LMR, SAA, FJQ, MM, BL and JMP designed the experiments. LMR, SAA, NR, RC, IM, TP, CCF, ADL and FJQ performed the experiments. LMR, SAA, FJQ, MM, BL and JMP analyzed the data. LMR and JMP wrote the paper. All the authors reviewed and approved the final manuscript. LMR and SAA contributed equally to this work.

## REFERENCES

Abualia R, Ötvös K, Novák O, Bouguyon E, Domanegg K, Krapp A, Nacry P, Gojon A, Lacombe B, Benková E (2022) Molecular framework integrating nitrate sensing in root and auxin-guided shoot adaptive responses. Proc Natl Acad Sci U S A 119: e2122460119

Aleman F, Nieves-Cordones M, Martinez V, Rubio F (2011) Root K(+) acquisition in plants: the Arabidopsis thaliana model. Plant and Cell Physiology 52: 1603–1612

Bañuelos MA, Garciadeblas B, Cubero B, Rodriguez-Navarro A (2002) Inventory and functional characterization of the HAK potassium transporters of rice. Plant Physiology 130: 784–795

Bañuelos MA, Klein RD, Alexander-Bowman SJ, Rodríguez-Navarro A (1995) A potassium transporter of the yeast Schwanniomyces occidentalis homologous to the Kup system of Escherichia coli has a high concentrative capacity. The EMBO Journal 14: 3021–3027

Barragan V, Leidi E, Andres Z, Rubio L, De Luca A, Fernandez J, Cubero B, Pardo J (2012) Ion Exchangers NHX1 and NHX2 Mediate Active Potassium Uptake into Vacuoles to Regulate Cell Turgor and Stomatal Function in Arabidopsis. The Plant Cell 24: 1127–1142

Beck G, Coman D, Herren E, Ruiz-Sola MA, Rodríguez-Concepción M, Gruissem W, Vranová E (2013) Characterization of the GGPP synthase gene family in Arabidopsis thaliana. Plant Mol Biol 82: 393–416

Bouguyon E, Brun F, Meynard D, Kubes M, Pervent M, Leran S, Lacombe B, Krouk G, Guiderdoni E, Zazimalova E, Hoyerova K, Nacry P, Gojon A (2015) Multiple mechanisms of nitrate sensing by Arabidopsis nitrate transceptor NRT1.1. Nat Plants 1: 15015

Chiu CC, Lin CS, Hsia AP, Su RC, Lin HL, Tsay YF (2004) Mutation of a nitrate transporter, AtNRT1:4, results in a reduced petiole nitrate content and altered leaf development. Plant Cell Physiol 45: 1139–1148

Corratgé-Faillie C, Lacombe B (2017) Substrate (un)specificity of Arabidopsis NRT1/PTR FAMILY (NPF) proteins. J Exp Bot 68: 3107–3113

Coskun D, Britto DT, Kronzucker HJ (2017) The nitrogen-potassium intersection: membranes, metabolism, and mechanism. Plant Cell Environ 40: 2029–2041

Cubero-Font P, Maierhofer T, Jaslan J, Rosales MA, Espartero J, Diaz-Rueda P, Muller HM, Hurter AL, Al-Rasheid KA, Marten I, Hedrich R, Colmenero-Flores JM, Geiger D (2016) Silent S-Type Anion Channel Subunit SLAH1 Gates SLAH3 Open for Chloride Root-to-Shoot Translocation. Curr Biol 26: 2213–2220

Deng K, Wang W, Feng L, Yin H, Xiong F, Ren M (2020) Target of rapamycin regulates potassium uptake in Arabidopsis and potato. Plant Physiol Biochem 155: 357–366

Dennison KL, Spalding EP (2000) Glutamate-gated calcium fluxes in Arabidopsis. Plant Physiol 124: 1511–1514

Dragwidge JM, Ford BA, Ashnest JR, Das P, Gendall AR (2018) Two Endosomal NHX-Type Na+/H+ Antiporters are Involved in Auxin-Mediated Development in Arabidopsis thaliana. Plant Cell Physiol 59: 1660–1669

Drechsler N, Zheng Y, Bohner A, Nobmann B, von Wiren N, Kunze R, Rausch C (2015) Nitrate-Dependent Control of Shoot K Homeostasis by the Nitrate Transporter1/Peptide Transporter Family Member NPF7.3/NRT1.5 and the Stelar K+ Outward Rectifier SKOR in Arabidopsis. Plant Physiol 169: 2832–2847

Duby G, Hosy E, Fizames C, Alcon C, Costa A, Sentenac H, Thibaud JB (2008) AtKC1, a conditionally targeted Shaker-type subunit, regulates the activity of plant K+ channels. Plant J 53: 115–123

Fan L, Zhao L, Hu W, Li W, Novak O, Strnad M, Simon S, Friml J, Shen J, Jiang L, Qiu QS (2018) NHX antiporters regulate the pH of endoplasmic reticulum and auxin-mediated development. Plant Cell Environ

Fang XZ, Liu XX, Zhu YX, Ye JY, Jin CW (2020) The K(+) and NO(3) (-) Interaction Mediated by NITRATE TRANSPORTER1.1 Ensures Better Plant Growth under K(+)-Limiting Conditions. Plant Physiol 184: 1900–1916

Fusco C, Guidotti E, Zervos AS (1999) In vivo construction of cDNA libraries for use in the yeast two-hybrid system. Yeast 15: 715–720

Gehl C, Waadt R, Kudla J, Mendel RR, Hänsch R (2009) New GATEWAY vectors for high throughput analyses of protein-protein interactions by bimolecular fluorescence complementation. Mol Plant 2: 1051–1058

Gierth M, Maser P, Schroeder JI (2005) The Potassium Transporter AtHAK5 Functions in K+ Deprivation-Induced High-Affinity K+ Uptake and AKT1 K+ Channel Contribution to K+ Uptake Kinetics in Arabidopsis Roots. Plant Physiol. 137: 1105–1114

Gietz RD, Schiestl RH (2007) Quick and easy yeast transformation using the LiAc/SS carrier DNA/PEG method. Nat Protoc 2: 35–37

Hedrich R, Geiger D (2017) Biology of SLAC1-type anion channels – from nutrient uptake to stomatal closure. New Phytologist 216: 46–61

Hirsch RE, Lewis BD, Spalding EP, Sussman MR (1998) A role for the AKT1 potassium channel in plant nutrition. Science 280: 918–921

Ho CH, Frommer WB (2014) Fluorescent sensors for activity and regulation of the nitrate transceptor CHL1/NRT1.1 and oligopeptide transporters. Elife 3: e01917

Ho CH, Lin SH, Hu HC, Tsay YF (2009) CHL1 functions as a nitrate sensor in plants. Cell 138: 1184–1194

Huang S, Maierhofer T, Hashimoto K, Xu X, Karimi SM, Müller H, Geringer MA, Wang Y, Kudla J, De Smet I, Hedrich R, Geiger D, Roelfsema MRG (2023) The CIPK23 protein kinase represses SLAC1-type anion channels in Arabidopsis guard cells and stimulates stomatal opening. New Phytol 238: 270–282

James GV, Patel V, Nordstrom KJ, Klasen JR, Salome PA, Weigel D, Schneeberger K (2013) User guide for mapping-by-sequencing in Arabidopsis. Genome Biol 14: R61

Jegla T, Busey G, Assmann SM (2018) Evolution and Structural Characteristics of Plant Voltage-Gated K(+) Channels. Plant Cell 30: 2898–2909

Jørgensen ME, Wulff N, Nafisi M, Xu D, Wang C, Lambertz SK, Belew ZM, Nour-Eldin HH (2017) Design and Direct Assembly of Synthesized Uracil-containing Non-clonal DNA Fragments into Vectors by USER(TM) Cloning. Bio Protoc 7: e2615

Jumper J, Evans R, Pritzel A, Green T, Figurnov M, Ronneberger O, Tunyasuvunakool K, Bates R, Žídek A, Potapenko A, Bridgland A, Meyer C, Kohl SAA, Ballard AJ, Cowie A, Romera-Paredes B, Nikolov S, Jain R, Adler J, Back T, Petersen S, Reiman D, Clancy E, Zielinski M, Steinegger M, Pacholska M, Berghammer T, Bodenstein S, Silver D, Vinyals O, Senior AW, Kavukcuoglu K, Kohli P, Hassabis D (2021) Highly accurate protein structure prediction with AlphaFold. Nature 596: 583–589

Jung JY, Shin R, Schachtman DP (2009) Ethylene mediates response and tolerance to potassium deprivation in Arabidopsis. Plant Cell 21: 607–621

Kato S, Hayashi M, Kitagawa M, Kajiura H, Maeda M, Kimura Y, Igarashi K, Kasahara M, Ishimizu T (2018) Degradation pathway of plant complex-type N-glycans: identification and characterization of a key α1,3-fucosidase from glycoside hydrolase family 29. Biochem J 475: 305–317

Katsiarimpa A, Anzenberger F, Schlager N, Neubert S, Hauser MT, Schwechheimer C, Isono E (2011) The Arabidopsis deubiquitinating enzyme AMSH3 interacts with ESCRT-III subunits and regulates their localization. Plant Cell 23: 3026–3040

Kopcsayová D, Vranová E (2019) Functional Gene Network of Prenyltransferases in Arabidopsis thaliana. Molecules 24

Krouk G, Lacombe B, Bielach A, Perrine-Walker F, Malinska K, Mounier E, Hoyerova K, Tillard P, Leon S, Ljung K, Zazimalova E, Benkova E, Nacry P, Gojon A (2010) Nitrate-regulated auxin transport by NRT1.1 defines a mechanism for nutrient sensing in plants. Dev Cell 18: 927–937

Lacombe B, Becker D, Hedrich R, DeSalle R, Hollmann M, Kwak JM, Schroeder JI, Le Novère N, Nam HG, Spalding EP, Tester M, Turano FJ, Chiu J, Coruzzi G (2001) The identity of plant glutamate receptors. Science 292: 1486–1487

Lacombe B, Thibaud JB (1998) Evidence for a multi-ion pore behavior in the plant potassium channel KAT1. J Membr Biol 166: 91–100

Léran S, Garg B, Boursiac Y, Corratgé-Faillie C, Brachet C, Tillard P, Gojon A, Lacombe B (2015) AtNPF5.5, a nitrate transporter affecting nitrogen accumulation in Arabidopsis embryo. Sci Rep 5: 7962

Lhamo D, Luan S (2021) Potential Networks of Nitrogen-Phosphorus-Potassium Channels and Transporters in Arabidopsis Roots at a Single Cell Resolution. Frontiers in Plant Science 12

Li H, Yu M, Du XQ, Wang ZF, Wu WH, Quintero FJ, Jin XH, Li HD, Wang Y (2017) NRT1.5/NPF7.3 Functions as a Proton-Coupled H(+)/K(+) Antiporter for K(+) Loading into the Xylem in Arabidopsis. Plant Cell 29: 2016–2026

Li J, Wu WH, Wang Y (2017) Potassium channel AKT1 is involved in the auxin-mediated root growth inhibition in Arabidopsis response to low K(+) stress. J Integr Plant Biol 59: 895–909

Li JY, Fu YL, Pike SM, Bao J, Tian W, Zhang Y, Chen CZ, Zhang Y, Li HM, Huang J, Li LG, Schroeder JI, Gassmann W, Gong JM (2010) The Arabidopsis nitrate transporter NRT1.8 functions in nitrate removal from the xylem sap and mediates cadmium tolerance. Plant Cell 22: 1633–1646

Lin SH, Kuo HF, Canivenc G, Lin CS, Lepetit M, Hsu PK, Tillard P, Lin HL, Wang YY, Tsai CB, Gojon A, Tsay YF (2008) Mutation of the Arabidopsis NRT1.5 nitrate transporter causes defective root-to-shoot nitrate transport. Plant Cell 20: 2514–2528

Maierhofer T, Scherzer S, Carpaneto A, Müller TD, Pardo JM, Hänelt I, Geiger D, Hedrich R (2024) Arabidopsis HAK5 under low K(+) availability operates as PMF powered high-affinity K(+) transporter. Nat Commun 15: 8558

Martín Y, Navarro FJ, Siverio JM (2008) Functional characterization of the Arabidopsis thaliana nitrate transporter CHL1 in the yeast Hansenula polymorpha. Plant Mol Biol 68: 215–224

Meng S, Peng JS, He YN, Zhang GB, Yi HY, Fu YL, Gong JM (2016) Arabidopsis NRT1.5 Mediates the Suppression of Nitrate Starvation-Induced Leaf Senescence by Modulating Foliar Potassium Level. Mol Plant 9: 461–470

Meyerhoff O, Müller K, Roelfsema MR, Latz A, Lacombe B, Hedrich R, Dietrich P, Becker D (2005) AtGLR3.4, a glutamate receptor channel-like gene is sensitive to touch and cold. Planta 222: 418–427

Morales de los Ríos L, Corratgé-Faillie C, Raddatz N, Mendoza I, Lindahl M, de Angeli A, Lacombe B, Quintero FJ, Pardo JM (2021) The Arabidopsis protein NPF6.2/NRT1.4 is a plasma membrane nitrate transporter and a target of protein kinase CIPK23. Plant Physiology and Biochemistry 168: 239–251

Mumberg D, Müller R, Funk M (1995) Yeast vectors for the controlled expression of heterologous proteins in different genetic backgrounds. Gene 156: 119–122

Nieves-Cordones M, Aleman F, Martinez V, Rubio F (2014) K(+) uptake in plant roots. The systems involved, their regulation and parallels in other organisms. J Plant Physiol 171: 688–695

Obrdlik P, El-Bakkoury M, Hamacher T, Cappellaro C, Vilarino C, Fleischer C, Ellerbrok H, Kamuzinzi R, Ledent V, Blaudez D, Sanders D, Revuelta JL, Boles E, Andre B, Frommer WB (2004) K+ channel interactions detected by a genetic system optimized for systematic studies of membrane protein interactions. Proc Natl Acad Sci U S A 101: 12242–12247

Okada K, Saito T, Nakagawa T, Kawamukai M, Kamiya Y (2000) Five geranylgeranyl diphosphate synthases expressed in different organs are localized into three subcellular compartments in Arabidopsis. Plant Physiol 122: 1045–1056

Pi L, Aichinger E, van der Graaff E, Llavata-Peris CI, Weijers D, Hennig L, Groot E, Laux T (2015) Organizer-Derived WOX5 Signal Maintains Root Columella Stem Cells through Chromatin-Mediated Repression of CDF4 Expression. Dev Cell 33: 576–588

Podar D, Maathuis FJM (2022) Primary nutrient sensors in plants. iScience 25: 104029

Raddatz N, Morales de Los Rios L, Lindahl M, Quintero FJ, Pardo JM (2020) Coordinated Transport of Nitrate, Potassium, and Sodium. Front Plant Sci 11: 247

Ragel P, Rodenas R, Garcia-Martin E, Andres Z, Villalta I, Nieves-Cordones M, Rivero RM, Martinez V, Pardo JM, Quintero FJ, Rubio F (2015) The CBL-Interacting Protein Kinase CIPK23 Regulates HAK5-Mediated High-Affinity K+ Uptake in Arabidopsis Roots. Plant Physiology 169: 2863–2873

Rissel D, Heym PP, Thor K, Brandt W, Wessjohann LA, Peiter E (2017) No Silver Bullet - Canonical Poly(ADP-Ribose) Polymerases (PARPs) Are No Universal Factors of Abiotic and Biotic Stress Resistance of Arabidopsis thaliana. Front Plant Sci 8: 59

Ródenas R, Vert G (2021) Regulation of Root Nutrient Transporters by CIPK23: ’One Kinase to Rule Them All’. Plant Cell Physiol 62: 553–563

Rodriguez-Navarro A, Ramos J (1984) Dual system for potassium transport in Saccharomyces cerevisiae. Journal of Bacteriology 159: 940–945

Rubio F, Fon M, Rodenas R, Nieves-Cordones M, Aleman F, Rivero RM, Martinez V (2014) A low K+ signal is required for functional high-affinity K+ uptake through HAK5 transporters. Physiol Plant 152: 558–570

Rubio F, Nieves-Cordones M, Aleman F, Martinez V (2008) Relative contribution of AtHAK5 and AtAKT1 to K+ uptake in the high-affinity range of concentrations. Physiologia Plantarum 134: 598–608

Run Y, Cheng X, Dou W, Dong Y, Zhang Y, Li B, Liu T, Xu H (2022) Wheat potassium transporter TaHAK13 mediates K(+) absorption and maintains potassium homeostasis under low potassium stress. Front Plant Sci 13: 1103235

Sanchez-Barrena MJ, Chaves-Sanjuan A, Raddatz N, Mendoza I, Cortes A, Gago F, Gonzalez-Rubio JM, Benavente JL, Quintero FJ, Pardo JM, Albert A (2020) Recognition and Activation of the Plant AKT1 Potassium Channel by the Kinase CIPK23. Plant Physiol 182: 2143–2153

Sena F, Kunze R (2023) The K(+) transporter NPF7.3/NRT1.5 and the proton pump AHA2 contribute to K(+) transport in Arabidopsis thaliana under K(+) and NO(3)(-) deficiency. Front Plant Sci 14: 1287843

Sentenac H, Bonneaud N, Minet M, Lacroute F, Salmon JM, Gaymard F, Grignon C (1992) Cloning and expression in yeast of a plant potassium ion transport system. Science 256: 663–665

Straub T, Ludewig U, Neuhauser B (2017) The Kinase CIPK23 Inhibits Ammonium Transport in Arabidopsis thaliana. Plant Cell 29: 409–422

Sun J, Bankston JR, Payandeh J, Hinds TR, Zagotta WN, Zheng N (2014) Crystal structure of the plant dual-affinity nitrate transporter NRT1.1. Nature 507: 73–77

Tang R-J, Zhao F-G, Yang Y, Wang C, Li K, Kleist TJ, Lemaux PG, Luan S (2020) A calcium signalling network activates vacuolar K+ remobilization to enable plant adaptation to low-K environments. Nature Plants 6: 384–393

Tascon I, Sousa JS, Corey RA, Mills DJ, Griwatz D, Aumuller N, Mikusevic V, Stansfeld PJ, Vonck J, Hanelt I (2020) Structural basis of proton-coupled potassium transport in the KUP family. Nat Commun 11: 626

Vanderauwera S, De Block M, Van de Steene N, van de Cotte B, Metzlaff M, Van Breusegem F (2007) Silencing of poly(ADP-ribose) polymerase in plants alters abiotic stress signal transduction. Proc Natl Acad Sci U S A 104: 15150–15155

Waadt R, Jawurek E, Hashimoto K, Li Y, Scholz M, Krebs M, Czap G, Hong-Hermesdorf A, Hippler M, Grill E, Kudla J, Schumacher K (2019) Modulation of ABA responses by the protein kinase WNK8. FEBS Lett 593: 339–351

Wang FZ, Zhang N, Guo YJ, Gong BQ, Li JF (2020) Split Nano luciferase complementation for probing protein-protein interactions in plant cells. J Integr Plant Biol 62: 1065–1079

Wang W, Hu B, Li A, Chu C (2020) NRT1.1s in plants: functions beyond nitrate transport. J Exp Bot 71: 4373–4379

Wang W, Yang X, Tangchaiburana S, Ndeh R, Markham JE, Tsegaye Y, Dunn TM, Wang GL, Bellizzi M, Parsons JF, Morrissey D, Bravo JE, Lynch DV, Xiao S (2008) An inositolphosphorylceramide synthase is involved in regulation of plant programmed cell death associated with defense in Arabidopsis. Plant Cell 20: 3163–3179

Wang YY, Cheng YH, Chen KE, Tsay YF (2018) Nitrate Transport, Signaling, and Use Efficiency. Annu Rev Plant Biol 69: 85–122

Xu J, Li HD, Chen LQ, Wang Y, Liu LL, He L, Wu WH (2006) A protein kinase, interacting with two calcineurin B-like proteins, regulates K+ transporter AKT1 in Arabidopsis. Cell 125: 1347–1360

Yang T, Feng H, Zhang S, Xiao H, Hu Q, Chen G, Xuan W, Moran N, Murphy A, Yu L, Xu G (2020) The Potassium Transporter OsHAK5 Alters Rice Architecture via ATP-Dependent Transmembrane Auxin Fluxes. Plant Communications 1: 100052

Yuan JS, Reed A, Chen F, Stewart CN, Jr. (2006) Statistical analysis of real-time PCR data. BMC Bioinformatics 7: 85

Zhang A, Ren HM, Tan YQ, Qi GN, Yao FY, Wu GL, Yang LW, Hussain J, Sun SJ, Wang YF (2016) S-type Anion Channels SLAC1 and SLAH3 Function as Essential Negative Regulators of Inward K+ Channels and Stomatal Opening in Arabidopsis. Plant Cell 28: 949–955

Zhang ML, Huang PP, Ji Y, Wang S, Wang SS, Li Z, Guo Y, Ding Z, Wu WH, Wang Y (2020) KUP9 maintains root meristem activity by regulating K(+) and auxin homeostasis in response to low K. EMBO Rep: e50164

Zhang S, Tajima H, Nambara E, Blumwald E, Bassil E (2020) Auxin Homeostasis and Distribution of the Auxin Efflux Carrier PIN2 Require Vacuolar NHX-Type Cation/H(+) Antiporter Activity. Plants (Basel) 9

Zhang X, Cui Y, Yu M, Su B, Gong W, Baluška F, Komis G, Šamaj J, Shan X, Lin J (2019) Phosphorylation-Mediated Dynamics of Nitrate Transceptor NRT1.1 Regulate Auxin Flux and Nitrate Signaling in Lateral Root Growth. Plant Physiol 181: 480–498

Zheng Y, Drechsler N, Rausch C, Kunze R (2016) The Arabidopsis nitrate transporter NPF7.3/NRT1.5 is involved in lateral root development under potassium deprivation. Plant Signal Behav 11: e1176819

